# Limitations of gene-dense datasets for inferring relationships in rapid radiations: An example from the Ice Plants

**DOI:** 10.64898/2026.11.18.767386

**Authors:** Oluwatomi E. Jacobs, Jordan Kressman, Shawn Arreguin, Tao Feng, Alfonso Timoneda, Caroline Edwards, Helene Tiley, Stephen A. Smith, Ya Yang, Samuel F. Brockington, Michael J. Moore, Joseph F. Walker

## Abstract

**Premise:** Aizoaceae (ice plants) are among the most rapidly radiating plant families on Earth, making them among the most difficult clades in the tree of life to resolve. Large, gene-rich datasets have been a powerful tool for exploring contentious relationships, and the Aizoaceae are an ideal model for testing the limits of gene-rich datasets for inferring the tree of life.

**Methods:** In this study, we assembled a phylotranscriptomic dataset comprising 28 newly sequenced and 11 publicly available transcriptomes to infer relationships within the Aizoaceae. We inferred gene trees from 4,845 putative orthologs extracted from 15,818 homologous gene families, which we also used to infer gene duplications. We tested whether gene-tree filtering approaches could identify gene subsets that resolve relationships within the clade.

**Results:** We found that, regardless of the gene-filtering approach or species-tree estimation, relationships among the Aizoaceae were largely congruent. Relationships within the subfamily Ruschioideae still showed little gene tree concordance. Our simulations show that inferring relationships within rapid radiations may be impossible because of short internodal branch lengths, regardless of the number of genes sampled. Gene duplication analysis at the base of the radiations provides candidates that potentially contributed to adaptations driving the radiation.

**Conclusions:** This study highlights that rapid radiations can be difficult to infer because of pervasive incomplete lineage sorting and noise, and that limited information within individual genes may make methods that rely on gene trees unsuitable for these relationships. This appears to be the case in Aizoaceae; however, these limitations suggest that synteny-based analyses will play an important role in future investigations of the tree of life.

## Introduction

The Aizoaceae are an iconic succulent plant family native to the arid regions of the Greater Cape Floristic Region of South Africa (**Manning & Goldblatt, 2012**). Despite the family’s abundance across South Africa, 28% of its species are data deficient according to the Red List of South African Plants. As such, the family is a top priority for taxonomic research in the country (**Victor & Smith, 2011**). To survive semi-arid desert conditions that many Aizoaceae grow in, members of the family exhibit a diverse range of life histories and striking morphological adaptations, making them an exemplar group for studying the extreme morphologies plants can exhibit (**Brockington et al., 2012**). Examples of such morphologies include ***Lithops***, the ‘living stones,’ a geophytic clade that produces just two leaves with translucent epidermal windows that channel sunlight to subterranean photosynthetic tissue (**Martin et al., 2013**). This window-leaf adaptation has evolved convergently in other Aizoaceae lineages, including ***Fenestraria***, in which club-shaped leaves remain mostly buried in sand, with only the translucent leaf tips exposed (**Martin et al., 2013**). Further morphological extremes are found in ***Conophytum***, notable for the extreme succulence of their fused leaf pairs and the frequently cryptic appearance of their vegetative bodies (**Opel, 2005**).

Much of this morphological diversity is associated with the family’s species richness, with five subfamilies (Aizooideae, Acrosanthoideae, Mesembryanthemoideae, Ruschioideae, and Sesuvioideae), 142 genera, and 1845 species (**Hernández-Ledesma et al., 2015**). Despite this species diversity, most Aizoaceae are endemic to southern Africa and have narrow geographic distributions (**Snijman, 2013**). The combination of high diversity and narrow distribution is an ecological indicator of rapid radiation, and researchers investigating the group have found the Aizoaceae to be among the fastest-diversifying plant family (**Klak et al., 2004; Valente et al., 2014**). As described by Klak et al. (2004), the rapid radiation of this group, specifically the subfamily Ruschioideae, was estimated to have occurred between 3.8 and 8.7 million years ago (mya), potentially coinciding with major climatic shifts in South Africa (5–10 mya). Three key innovations are hypothesized to have facilitated the clade’s diversification during this climatic transition from a moist to an extremely dry environment: (1) wide-band tracheids, which are specialized to prevent the collapse of the primary wall and are strongly associated with adaptations to withstand water stress; (2) flattened, cylindrical/trigonous leaf shapes that reduce water loss in dry conditions; (3) core-specialized hygrochastic capsules that open only when moistened to release a portion of their seeds in the rain. As with any rapidly radiating group, the relationships among the Aizoaceae, particularly within the Ruschioideae, have proven a near-impossible challenge for phylogenetics.

To investigate relationships within this particularly diverse group, previous molecular phylogenetic studies have used nuclear ribosomal DNA to examine the tribe Sesuvioideae (**Hassan et al., 2005**) and plastid markers (**Klak et al., 2003**; **Thiede et al., 2007**). More efficient bioinformatics pipelines and sequencing techniques have enabled more gene- and taxon-rich investigations. As such, recent systematics work has largely consisted of studies leveraging the plastome data (**Klak et al., 2013**; **Liede-Schumann et al., 2020**; **Klak et al., 2024**). For the subfamily Sesuvioideae, plastid and ribosomal DNA markers have helped reveal a preliminary timeline of the development of C_4_ photosynthesis within the subfamily (**Bohley et al., 2015**) and, with genome skimming, the group’s taxonomy and challenges, such as conflicting gene tree signals, are being uncovered (**Kadereit et al., 2026**).

Despite these efforts, much of our current knowledge of Aizoaceae systematics is marked by the challenges posed by this group and the polytomies recovered within the Ruschioideae. Exploring these patterns with a large, nuclear dataset can help us better understand these contentious relationships. Phylotranscriptomics is a cost-effective approach to obtaining data for thousands of genes (**Yang et al., 2017; Arreguin et al., 2025**), and phylotranscriptomic studies have shed light on relationships across the plant tree of life (**Walker et al., 2018**; **One Thousand Plant Transcriptomes Initiative, 2019**). To date, no phylotranscriptomic study has focused on Aizoaceae. Gene-rich datasets provide broad genome sampling that can help elucidate contentious relationships and the molecular evolutionary processes underlying them. These processes often arise at the microevolutionary scale; however, they can be inferred at the macroevolutionary scale, helping to bridge the two. Such evolutionary processes that can be uncovered by phylotranscriptomics include gene and genome duplication, gene tree conflict, and changes in rates of molecular evolution (**Yang et al., 2015**).

In this study, we aim to leverage phylotranscriptomics to resolve the relationships within the Aizoaceae, with a focus on the Ruschioideae. We use machine-learning-based phylogenetic difficulty prediction and an information-based approach to filter gene trees for the optimal genes and to test whether gene filtering may assist in phylogenetic inference. Finally, to further our understanding of what may have turned the Ruschioideae into a systematics challenge and promoted the greatest known radiation in the plant tree of life, we investigate gene duplications predicted to have occurred at the base of the group. This study provides insights into the difficulties encountered in studying radiation and the molecular bases that may contribute to them.

## Materials and Methods

### Taxon sampling and data acquisition

Plants for the 28 newly sequenced Aizoaceae samples were predominantly collected from living collections (**Supplementary Table 1**). Taxa from three of the five recognized subfamilies, Ruschioideae, Sesuvioideae, and Mesembryanthemoideae (**Klak et al., 2017**) were sampled. Within Ruschioideae, two of the four tribes, Ruschiae and Drosanthemeae (**Liede-Schumann et al., 2020**), were sampled and within Sesuvioideae, one of the two tribes, Sesuvieae, was sampled (**Sukhorukov et al., 2018**). 18 species from Rushioideae, the fastest radiating tribe, were sampled. These include *Fenestraria aurantiaca, Glottiphyllum uncatum, Hallianthus planus, Juttadinteria kovisimontana, Trichodiadema cf. marlothii, Stoeberia utilis, Ruschianthus falcatus, Pleiospilos magnipunctatus, Leipoldtia lunata, Monilaria moniliformis, Faucaria felina, Acrodon bellidiflorus, Juttadinteria attenuata, Titanopsis hugo-schlechteri, Argyroderma fissum, Cephalophyllum cupreum, Gibbaeum album, Dracophilus dealbatus.* Tissues were flash-frozen in liquid nitrogen, then stored at -80 °C until extraction. RNA was extracted using PureLink Plant RNA Reagent (Invitrogen, Carlsbad, CA, USA) followed by treatment with the TURBO DNA-free kit (Ambion, Carlsbad, CA, USA) to remove genomic DNA. RNA from *Glottiphyllum uncatum* and *Pleiospilos magnipunctatus* was extracted using an initial hot acid phenol extraction step, followed by purification with the QIAGEN RNeasy Plant Mini Kit (Hilden, Germany) and on-column removal of genomic DNA using the QIAGEN RNase-Free DNase Set (Hilden, Germany). For the initial hot acid phenol extraction, less than 100 mg of ground plant tissue was mixed with 0.9 mL of LiCl extraction buffer (0.1M Tris pH9, 1% SDS, 0.1M LiCl, 0.01M EDTA pH8), followed by the addition of 0.9 mL of saturated acid phenol, both preheated to 60 °C. Samples were shaken vigorously and centrifuged at 10,000 rpm for 10 min. The upper aqueous phase was recovered, mixed vigorously with 0.9 mL of chloroform:isoamyl alcohol (24:1), and centrifuged again at 10,000 rpm for 10 min. The resulting supernatant was recovered and mixed by inversion with an equal volume of isopropanol. Samples were incubated at 4 °C for 1 hour and centrifuged at 13,000 rpm for 10 min. The resulting pellets were washed with chilled 70% ethanol and centrifuged again, twice. Pellets were allowed to dry, and resuspended in 200 µL of DEP-DI-water, mixed with 200 µL of 4 M LiCl, and incubated overnight at 4 °C. Samples were then centrifuged at 13,000 rpm for 30 min, and the supernatant was discarded. The resulting pellets were washed three times with 70% ethanol, with centrifugation at 13,000 rpm for 2 min after each wash. Finally, the pellets were dried and resuspended in 30–50 µL of DEP-DI water.

Sequencing was performed by GeneWiz on an Illumina HiSeq platform, with library prep using polyA selection in a 2x150 bp configuration, generating between 10,743,540 and 53,478,932 raw paired-end reads (**Supplementary Table 2**). We downloaded reads for an additional six Aizoaceae samples from the National Center for Biotechnology Information Sequence Read Archive (NCBI SRA) and five outgroups (*Anisomeria littoralis*, *Gisekia pharnaceoides*, *Salpianthus purpurascens*, *Rivina humilis* and *Kewa bowkeriana*), spanning a grade within which Aizoaceae is nested based on previous studies (**Walker et al., 2018**; **One Thousand Plant Transcriptomes Initiative, 2019**).

### Transcriptome Assembly and Quality Assessment

The individual transcriptomes were uniformly assembled from raw reads using the Semblans v1.0.2 bioinformatics pipeline (**Woodcock-Girard et al., 2025**), which in short consists of the following preprocessing steps: (1) the paired-end RNA-seq reads are corrected for base pair errors using Rcorrector v1.0.7 (**Song and Florea, 2015**); (2) The adapters are removed and sequences are quality trimmed using Trimmomatic v0.39 (**Bolger et al., 2014**); (3) Non-plant contamination is identified and removed using Kraken2 v2.1.2 (**Wood et al., 2019**) with removal based on the archaea, bacteria, viral, fungi, human, protozoa databases (4) Over-represented reads are identified using FastQC (**Andrews, 2010**) and filtered to generate the “cleaned reads”. The cleaned reads are then assembled using the de novo assembler Trinity v2.15.1 (**Grabherr et al., 2011**). The postprocessing of the assembled isoforms follows the procedure of: (1) Chimeric sequences are identified and removed using the Semblans plant database (https://github.com/gladshire/Semblans), with the Diamond v2.1.7 (**Buchfink et al., 2015**) implementation of the BLASTX (**Camacho et al., 2009**) algorithm, following the **Yang and Smith (2013)** approach; (2) The remaining sequences are collapsed to predict gene-level representation using Corset (**Davidson & Oshlack, 2014**), with the mapping performed using Salmon v1.10.0. The final transcripts are then converted to the best-supported open reading frames using TransDecoder v5.7.0, with homology identification conducted using the same plant database as chimera detection and the BLASTP process as implemented in Diamond v.2.1.7.

The final inferred ORFs were annotated with PANTHER v2.2 (**Mi et al., 2021**), which uses HMMER v3.4 (**Finn et al., 2011**) to detect sequence similarity. The assembled transcriptomes were assessed for completeness using Benchmarking Universal Single-Copy Orthologs v5.7.1 (BUSCO) (**Simão et al., 2015**) with the **eudicots_odb10** dataset (n = 2,326 BUSCOs) and the **-- mode euk_tran setting**. The quality of the assembled contigs was further evaluated with TransRate2 (https://github.com/ericbretz/transrate2), as utilized in **Tyszka et al. (2026)**, which is an updated approach to assess read support, coverage uniformity, and structural consistency of assembled transcripts by mapping reads back to the assemblies, using the metrics outlined in **Smith-Unna et al. (2016)**.

### Homology estimation

Homology estimation was conducted following the procedure developed by **Yang and Smith (2014)** and further refined by **Morales-Briones et al. (2021)**. Each inferred transcriptome was first compressed to remove redundancy using CD-HIT v.4.8.1 with parameters -c 0.99 -n 10 -r 0. An all-by-all similarity search was conducted using BLASTN with the settings “--evalue 10 – max_target_seqs 1000” across all transcriptome assemblies to infer putative homolog groups. These groups were then refined using Markov Clustering mcl v14-137 (**Van Dongen, 2000**) with an inflation value of 1.4, selected as a moderate clustering stringency that balances cluster fragmentation and over-merging. The resulting homolog clusters were used as input for the subsequent tree-based homology refinement.

Tree-based homolog refinement was then conducted on each resulting putative homolog. First, alignments were generated using MAFFT v7.490 (**Katoh and Standley, 2013**) with the settings “--auto --maxiterate 1000”. The resulting alignments were then cleaned to a minimum of 10% column occupancy using the Phyx v.1.2 program *pxclsq* (**Brown et al., 2017**). Phylogenetic trees were inferred using maximum-likelihood as implemented in IQ-TREE2 v2.3.6 (**Minh et al., 2020**) under the GTR+G model of evolution. Any branches on the resulting homolog trees with a relative value of 0.1 subs/site or 10 times longer than the sister branch, or an absolute value of 0.2 subs/site were removed. Any clade consisting of only the same taxa was then masked, so only the most character-rich tip in the cleaned alignment remained. Finally, any clade that contained at least 10 taxa and was on an internal branch of 0.2 subs/site or greater was split into a separate homolog. All resulting homologs were then evaluated to determine whether the tree-based refined homolog had changed in the number of sequences. All homologs that changed in sequence number were then subjected to the tree refinement procedure until no further changes occurred. Once no further changes were detected, final homolog trees were rerun with 1000 ultra bootstrap replicates (**Hoang et al., 2018**) for support. As no recurrent genome duplications have previously been reported in Aizoaceae, orthologs were extracted using Maximum Inclusion (MI). The MI method identifies the largest monophyletic subtree within each homolog tree that contains at most one sequence per taxon, thereby extracting single-copy orthologs while maximizing taxon occupancy across the given dataset. The relative and absolute branch length cutoffs were set to 0.1 and 0.2 subs/site, respectively, and a minimum taxon occupancy filter of 20 taxa were used in the ortholog extracting step.

### Gene and species tree estimation

We aligned the nucleotide sequences for the orthologs extracted in the homolog estimation step using PRANK v.170427 (**Löytynoja, 2013**). We trimmed the resulting alignments to a minimum of 30% column occupancy using *pxclsq*, and inferred gene trees using maximum likelihood in IQ-TREE2 v2.3.6 under the GTR+G model of evolution, with 1000 ultrafast bootstrap replicates for support. Species tree estimation was conducted using two approaches: (1) a concatenation-based approach, where all ortholog alignments were concatenated into a partitioned supermatrix using the Phyx program *pxcat* (**Brown et al., 2017**) and a tree was inferred using maximum-likelihood as implemented in IQ-TREE2 v2.3.6, with 1000 ultrafast bootstrap replicates for support; (2) a coalescent-based approach using the maximum quartet support species tree algorithm as implemented in ASTRAL v5.7.8 (**Zhang et al., 2018**) with all inferred orthologs as input and Local Posterior Probabilities as support (**Sayyari and Mirarab, 2016**). Gene tree conflict was estimated by first rooting the trees on the outgroups with the Phyx program *pxrr* (**Brown et al., 2017**), resulting in 2857 gene trees that could be rooted on the outgroups, and the gene trees were mapped to the supermatrix tree using PhyParts v0.0.1 (**Smith et al., 2015**) with a 95% bootstrap filter.

### Tree Certainty Score Calculations

To calculate information-theoretic branch support scores, we generated 200 regular bootstrap replicates for each ortholog gene tree using IQ-TREE2 v2.3.6 under the GTR+G model of nucleotide evolution. The bootstrap tree topologies served as the basis for calculating the Relative Tree Certainty All (TCA) score (**Salichos et al., 2014**). The TCA score is an information-theoretic metric that quantifies the overall degree of conflict among trees in a given tree set. It is the sum of all Internode Certainty All Scores (ICA) scores divided by the total number of internodes. The ICA scores are calculated by taking a bipartition in the tree of interest (the gene tree), identifying all bipartitions in a tree set (the bootstrap trees) that intersect with the bipartition of interest, and inferring how similar the bipartitions are to both the bipartition of interest and to one another. If the bipartition of interest is identical to all intersecting bipartitions in the tree of interest, the ICA score is 1.0. If all intersecting bipartitions are identical to one another but different from the bipartition of interest, the ICA score is -1.0. Thus, the closer the ICA score is to 0, the greater the differences among the intersecting bipartitions. Since the TCA score is directly related to the ICA score, a TCA score of 1.0 would indicate complete uniformity across the bootstrap tree set and the gene tree. The TCA score for all 4845 genes using RAxML v8.2.12 (**Stamatakis, 2014**), with a custom script to batch-process the results (https://github.com/tomi-jacobs/InfoCheck).

### Pythia score calculations

As another approach to evaluating gene-by-gene predicted difficulty of phylogenetic inference for each ortholog alignment, we used the multiple-sequence alignment evaluation program Pythia2.0 (**Haag and Stamatakis, 2025**). Pythia uses RAxML-NG v1.2.2 (**Kozlov et al., 2019**). It employs a Gradient Boosted Trees regressor to predict the likelihood that multiple independent maximum-likelihood tree inferences will converge on a single topology. In short, Pythia provides a score that indicates how difficult it will be to identify a single maximum-likelihood topology from a sequence alignment. The score ranges from 0.0 to 1.0, with 0.0 indicating that predicting a single tree from the alignment is easy and 1.0 indicating that inferring a single tree is difficult. A custom script was used to automate the process across all 4845 alignment files (https://github.com/tomi-jacobs/InfoCheck).

### Gene Tree Filtering Concordance Analysis

The TCA and Pythia scores were used to filter the 4,845 orthologs (ALL dataset) into three additional datasets: (1) Genes with a TCA score ≥ 0.3 were retained (BIPART [n = 450]); (2) Genes with a Pythia score ≤ 0.3 were retained (PYTH [n = 2092]); (3) Genes meeting both criteria (Pythia score ≤ 0.3 ≤ TCA score) (BOTH [n = 432]). Genes with a TCA score ≥ 0.3 were retained (BIPART [n=450]), representing gene trees with concordance exceeding conflict across bipartitions. Genes with a Pythia score ≤ 0.3 were retained (PYTH [n=2092]), representing the “easy” classification threshold below which independent maximum likelihood inferences predicted reliably to converge on a single topology. Both thresholds were then selected to represent a third and conservative filter that would retain only gene trees that meet both criteria. For each dataset, a species tree was inferred using ASTRAL and IQ-TREE2. Gene concordance factors (**Minh et al., 2020a**) for each dataset were estimated using IQ-TREE2, with each set analyzed on its respective tree.

### Simulating idealized gene tree sets

We simulated four datasets of 4,850 nucleotide sequences using seq-gen v1.3.4 (**Rambaut and Grass, 1997)**, rounded up from the 4,845 orthologs. Each dataset contained sequences of four lengths (500 bp, 1000 bp, 2000 bp, and 5000 bp). Simulations were carried out under the GTR model of nucleotide evolution with gamma rate variation divided into four categories, using the rooted ALL supermatrix tree as the input tree. For each dataset, we inferred gene trees for the simulated nucleotide sequences using IQ-TREE2 with the GTR+G model of evolution. Gene concordance factors were calculated using IQ-TREE2 on the ALL supermatrix topology for trees inferred for each simulated dataset.

### Plastome Inference

To construct the plastome dataset from transcriptomes, total raw reads for each sample were first cleaned using the preprocess function in Semblans v1.0.1, which ended with the Trimmomatic step. We then used HybPiper v2.3.3 (**Johnson et al., 2016**) to map and assemble reads. Mapping was conducted using a custom nucleotide-based target file for the plastome, derived from publicly available plastomes (**Supplementary Table 3**). Genes in the target file were present in at least four genomes; if two copies existed, either the longest one was kept or, if they were of equal length, the first that appeared in the file. Assembly of the reads was conducted by mapping them to the constructed target files using BWA v0.7.19 (**Li and Durbin, 2009**). Reads were then assembled using SPAdes v4.2.0 (**Prjibelski et al., 2020**), and coding sequences were extracted using Exonerate v2.4.0 (**Slater & Birney, 2005**)

To infer the plastome phylogeny, respective loci were aligned using PRANK with default settings. Alignments were cleaned for a minimum of 10% column occupancy using *pxclsq*. Cleaned alignments were then concatenated into a supermatrix using *pxcat*. The supermatrix was then visually inspected for misalignment. If sequences showed obvious signs of misalignment, they were removed, and loci were then realigned, cleaned, and concatenated into a new supermatrix, following the above procedure. Lastly, a maximum-likelihood tree was inferred using IQ-TREE2 v2.0.7 with linked rate variation among partitions and 1000 ultrafast bootstrap replicates for support.

### Detection of contamination and mislabeling

Two approaches were used to identify potential contamination and mislabeling. The first was to download *matK* and *rbcL* sequences of Aizoaceae from NCBI and test whether our samples are sister to those sequences in a phylogeny. We aligned existing and newly generated *matK* and *rbcL* sequences independently using PRANK (v.170427) with default settings. Alignments were then cleaned for a minimum of 10% column occupancy using *pxclsq*. IQ-TREE2 was then used to infer a maximum likelihood tree using the GTR+G substitution model with 1000 ultrafast bootstrap replicates for support.

The second approach was to examine whether HybPiper predicted multiple contigs for plastome genes. HybPiper is more likely to predict multiple copies of genes such as *matK, rbcL, and ndhF,* if sampled transcriptomes are cross-contaminated. To avoid inflated numbers due to small fragments, we applied a minimal length cutoff of 800 base pairs. One limitation of any approach to identify potential spillover that cannot be overcome is the high similarity among Ruschioideae species, which in theory could lead to near-identical plastomes being predicted from multiple species. More discussion on the similarity can be found below.

### Gene duplication analysis

Rooted clades with a minimum of four taxa containing ingroup-specific gene duplications were extracted from the final 15,818 homolog trees using the extract_clades.py script (**Yang et al., 2015**). The extraction yielded 13,822 rooted clades, which were mapped to the species tree using PhyParts v0.0.1 (**Smith et al., 2015**), with a 95% filter and the verbose setting enabled to identify rooted clades that duplicated at specific locations. Functional annotations were then assigned to each extracted clade by identifying the PANTHER annotation that matches the sequences within the clade.

## Results

### Assembly quality

We assessed assembly quality using Benchmarking Universal Single Copy Orthologs (BUSCOs) and TransRate scores for each assembly. The BUSCO scores ranged from 32.3% complete BUSCOs for *Sesuvium verrucosum* to 89.6% complete BUSCOs for the outgroup *Salpianthus purpurascens,* as shown in **Supplementary Figure 1**. The TransRate scores indicated generally high sequence accuracy across assemblies. The contig scores, an assessment of the reliability of the contigs, ranged from 0.08 to 0.51 with individual component scores ranging as follows: s(C_nuc_), an assessment of similarity in base pairs between reads and contigs, ranged from 0.68 to 1.00; s(C_cov_), the proportion of nucleotides within a contig that have read support, ranged from 0.31 to 0.93; s(C_ord_), the proportion of reads that map in the proper orientation, ranged from 0.44 to 0.86; and s(C_seg_), the proportion of reads without unexpected segmentation arising from chimeric assemblies, ranged from 0.55 to 1.00 across taxa (**Supplementary Figure 2**). The sCnuc was consistently the highest-scoring component, with most assemblies achieving near-perfect nucleotide accuracy.

For the contamination analysis, we obtained 20 *matK* samples from NCBI, and of those 20 possible comparisons, seven samples ended up sister to another species (**Supplementary Figure 3).** However, none of these comparisons were well supported (UFBoot <= 95) and all had short branch lengths (subs/bp < 0.05). *Sesuvium portulacastrum* had the highest support for not being sister to its corresponding NCBI sequence (UFBoot = 86) but was not obtained from this study. Therefore, this sample could not have been contaminated by sequences in this study. For *rbcL,* we obtained 13 sequences spanning 12 species. We found similar results, with seven taxa not sister to their GenBank counterparts, and none were well supported. Three of these, *S. portulacastrum, Delosperma echinatum* and *Stoeberia utilis*, were also in the *matK* tree. When examining how many contigs were predicted for *matK*, *rbcL,* and *ndhF*, we recovered only one contig in each case (**Supplementary Table 4**).

### Homology Estimation and Ortholog Extraction

We applied the **Yang and Smith (2014)** approach to tree-based homolog cleaning to the initial 21,466 homologs inferred from BLAST and MCL-based clustering. Cleaning for each homolog was performed until the number of sequences in the homolog after cleaning matched the number before cleaning, indicating that, given the specified parameters, the tree no longer benefited from further cleaning. As a result, most gene trees needed at least one cleaning, with a maximum of 20 iterations; however, a steep drop-off was noticeable after the first round of cleaning (**Supplementary Figure 4**). In total, the cleaning resulted in 15,818 homolog clusters, of which 4,845 ortholog clusters contained at least 20 taxa, and 764 had complete taxon occupancy. Based on gene filtering of the ALL dataset (**Figure 1A)**, the PYTH dataset showed a noticeable increase in gene length, with a greater increase for the BIPART dataset and a slight decrease relative to the BIPART dataset for the BOTH dataset (**Figure 1B)**.

**Figure 1.**
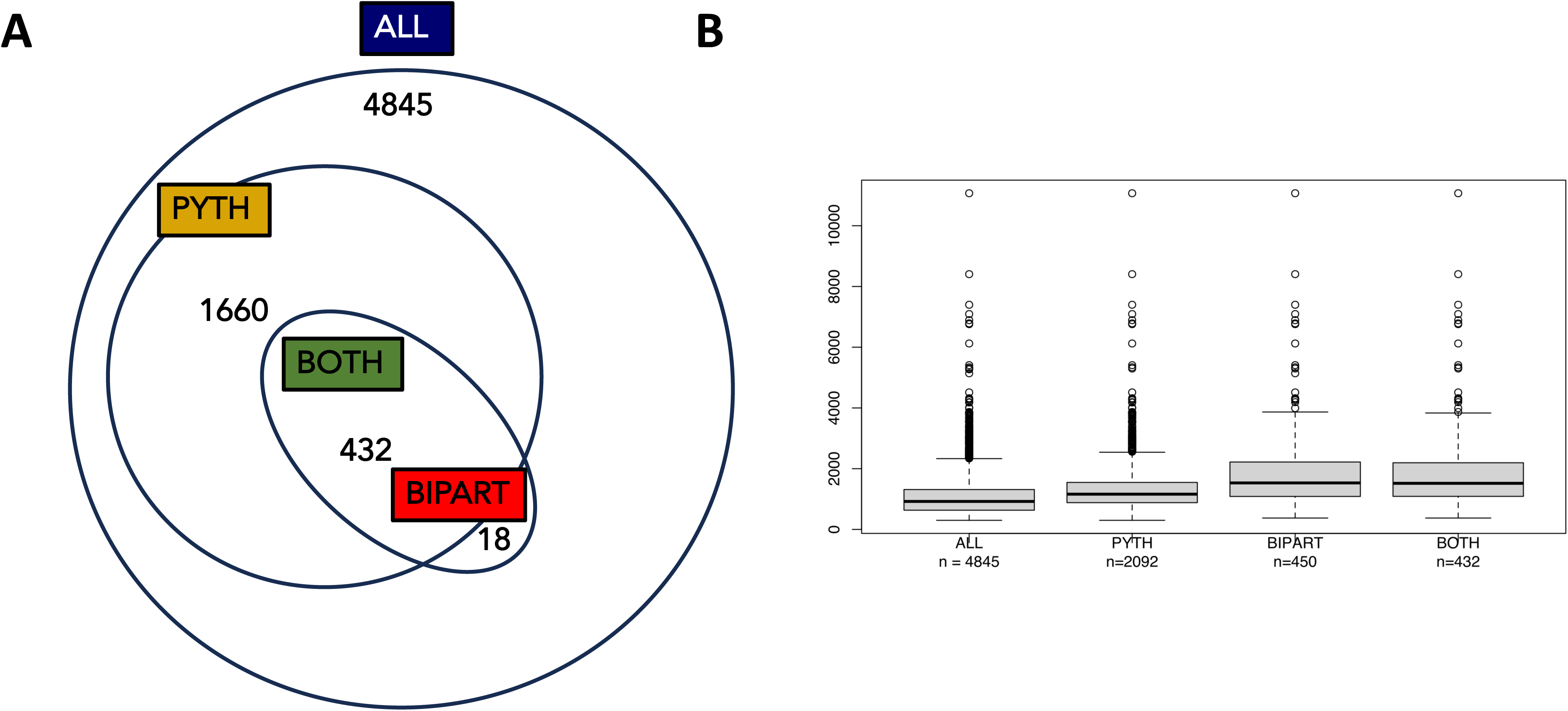
Number of sequences in each dataset and length of the sequences within the datasets. A) Total number of orthologs in the study for each filtered dataset. B) Box plots of the length of sequences remaining after each filtering approach.

### Inferred Species Relationships

Regardless of the tree inference method or filtration approach used, the inferred species relationships remained largely stable, and the Aizoaceae were recovered as monophyletic (**Supplementary Figure 5**). The three sampled subfamilies were recovered as reciprocally monophyletic, with the Sesuvioideae sister to Mesembryanthemoideae and Ruschioideae. Within Sesuvioideae, *Trianthema portulacastrum* was sister to a clade containing *Zaleya pentandra* and the monophyletic *Sesuvium*.

The Ruschioideae are among the groups whose relationships vary with the tree inference method and the filtering approach. As such, the description will follow the ALL supermatrix tree’s inferred relationships, with differences among datasets described in the next section. The earliest divergence, *Drosanthemum candens* is recovered as sister to the rest of Rushieae.

The next clade within the grade is *Delosperma echinatum*, *Trichodiadema pomeridianum*, and *Trichodiadema* cf. *marlothii* form a clade, with *Glottiphyllum uncatum* sister to that group, and *Gibbaeum album* sister to all three. This entire group forms one of the major lineages within core Ruschioideae (**Figure 2**). The sister to the remaining ingroup is a clade that contains *Monilaria moniliformis,* sister to a clade with *Pleiospilos magnipunctatus* sister to *Rhombophyllum nelii*, *Acrodon bellidiflorus*, and *Faucaria felina*. The next clade includes *Conophytum uviforme* and *Titanopsis hugo-schlechteri* as the sister taxon to *Juttadinteria attenuata, Juttadinteria kovisimontana*, *Dracophilus dealbatus, Psammophora modesta*, and *Ruschianthus falcatus*. Sister to the rest of the ingroup is a clade consisting of *Argyroderma fissum* sister to a clade of *Stoeberia utilis, Stoeberia carpii* as sisters, then *Eberlanzia ebracteata*, then *Fenestraria rhopalophylla* and *Fenestraria aurantiaca,* then *Cephalophyllum cupreum* sister to *Leipoldtia lunata* and *Hallianthus planus*.

**Figure 2.**
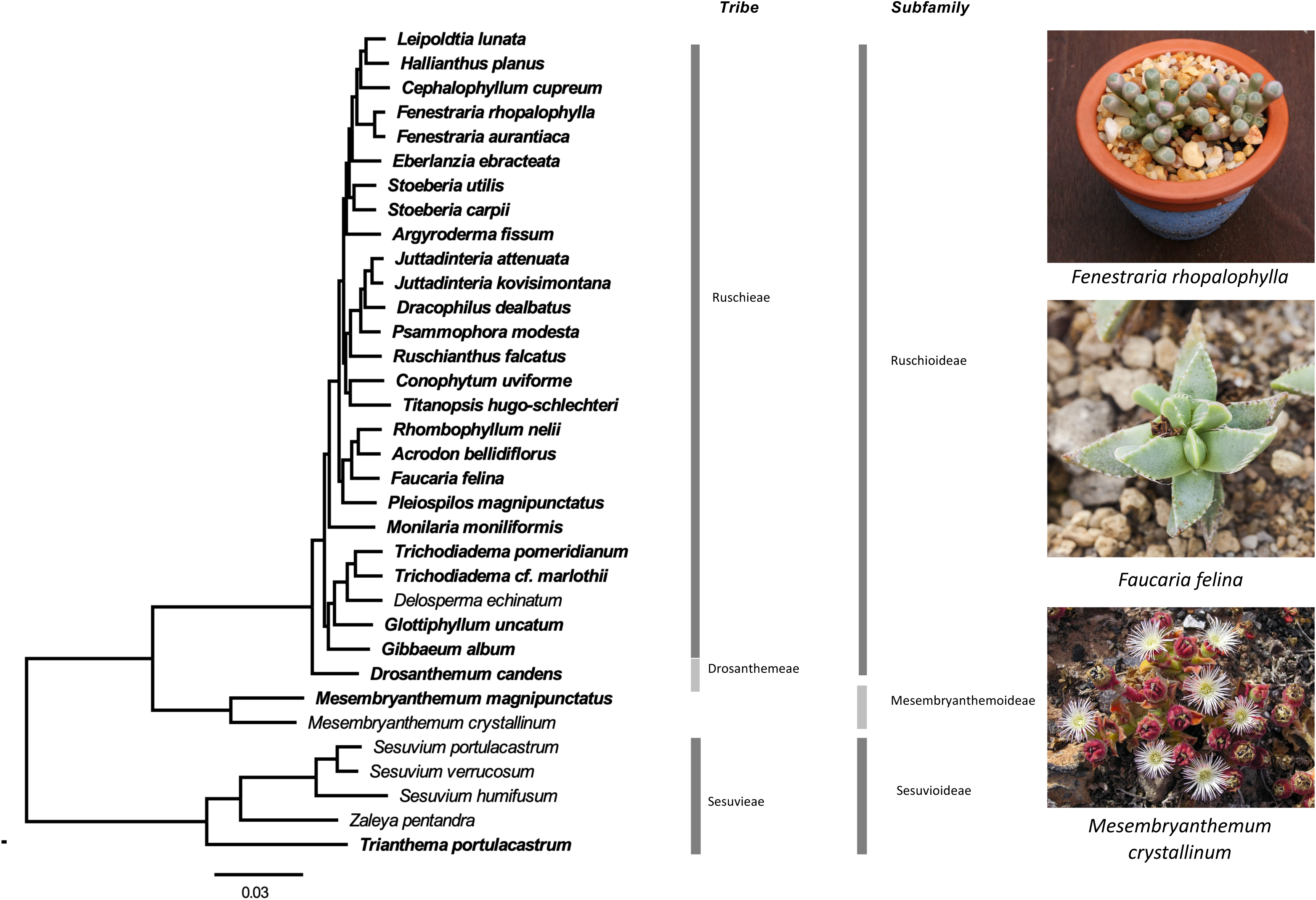
The relationships inferred using the ALL dataset and the supermatrix tree approach. All nodes have 100% ultrafast bootstrap support. Taxa newly sequenced for the study are bolded. Outgroups have been removed. Photo credits: *Fenestrania rhopalophylla* (author: Dr. David Midgley; license: CC BY-SA 2.5), *Faucaria felina* (author: Diego Delso; license: CC BY-SA 3.0), *Mesembryanthemum crystallinum* (author: Yummifruitbat; license: CC BY-SA 4.0).

### Concordance among species tree inference methods and data filters

Across datasets and methods, four unique species relationships were inferred, with conflicts between methods occurring at short internodes within the Ruschioideae (**Figure 3A; Supplementary Figure 6)**. Across all eight inferred species trees, thirty-one of the thirty-six edges were concordant (**Figure 3A**). In the ALL and PYTH datasets, the supermatrix and ASTRAL topologies were identical, except for the position of *Titanopsis hugo-schlechteri* (**Supplementary Figure 5**). In the supermatrix, *Titanopsis hugo-schlechteri* was sister to *Conophytum uviforme* (ALL gCF = 8.5%; PYTH gCF = 9.49%) and formed a clade sister to [*Juttadinteria attenuata, Juttadinteria kovisimontana, Dracophilus dealbatus, Psammophora modesta, Ruschianthus falcatus*] (ALL gCF = 1.13%; PYTH gCF = 1.86%), whereas in the ASTRAL tree, *Titanopsis hugo-schlechteri* was sister to a clade containing *[Leipoldtia lunata, Hallianthus planus, Cephalophyllum cupreum, Fenestraria rhopalophylla, Fenestraria aurantiaca, Eberlanzia ebracteata, Stoeberia carpii, Stoeberia utilis, Argyroderma fissum*] (ALL gCF = 1.13%; PYTH gCF = 4.25%). The relationship of *Conophytum uviforme* sister to [*Juttadinteria attenuata, Juttadinteria kovisimontana, Dracophilus dealbatus, Psammophora modesta, Ruschianthus falcatus*] had (ALL gCF = 3.08%; PYTH gCF = 7.64%). These relationships were supported by 100% UFBoot and 70% UFBoot for the ALL and PYTH supermatrices, respectively, and 1.0 LPP for both the ALL and PYTH ASTRAL trees.

**Figure 3.**
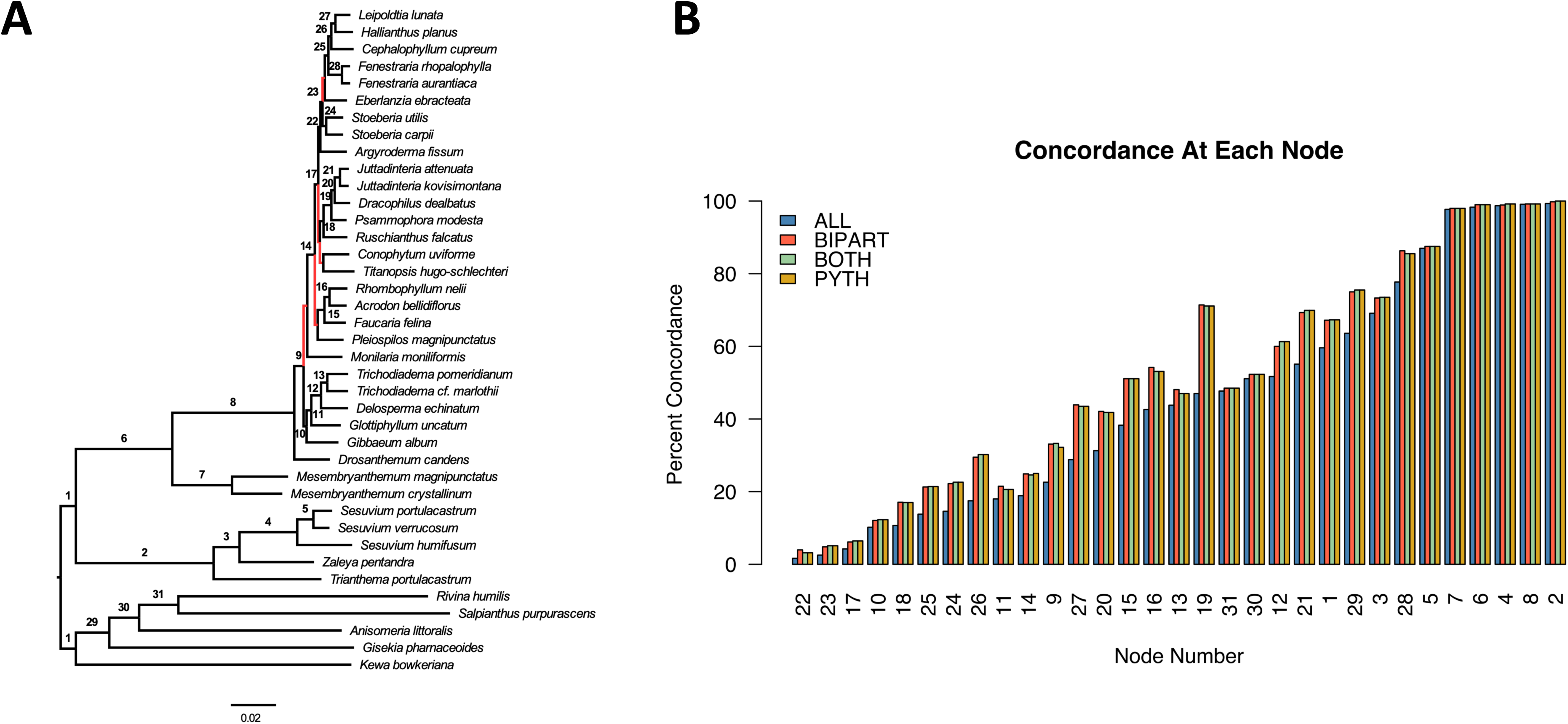
Gene tree concordance is low across relationships within the Ruschiodeae. A) The tree inferred from a supermatrix using the ALL dataset, with all branches that were not concordant across tree inference and datasets labeled in red. Node labels correspond to the concordance graph in B. B) Graph of the percent of gene trees concordant at each node for each dataset.

In the BIPART and BOTH trees, the supermatrix topology placed *Eberlanzia ebracteata* as the sister to [*Stoeberia utilis* and *Stoeberia carpii*] (BIPART gCF = 8.85%; BOTH gCF = 9.34%), with ultrafast bootstrap support of 77% and 70%, respectively. The ASTRAL tree placed *Eberlanzia ebracteata* as a sister to a clade of [*Leipoldtia lunata, Hallianthus planus, Cephalophyllum cupreum, Fenestraria aurantiaca, Fenestraria rhopalophylla]* (BIPART gCF = 5.25%; BOTH gCF = 5.54%) with Local Posterior Probability support of 0.49 and 0.52, respectively.

### Gene tree concordance across datasets

Across the thirty-one shared edges, the ALL dataset consistently had the lowest percentage of concordant genes, with the edge supporting (*Psammophora modesta, Dracophilus dealbatus, Juttadinteria attenuata, Juttadinteria kovisimontana*) showing the greatest difference between the unfiltered (ALL gCF = 47%) and the filtered datasets (BIPART gCF = 71.4%; BOTH gCF = 71.1%; PYTH gCF = 71.1%) (**Figure 3B**). The Ruschioideae were monophyletic with at least 99% gene concordance across all datasets. However, all conflicts among datasets occurred within the Ruschioideae, and inter-subfamilial relationships were recovered with low gene tree concordance.

### Simulations using different sequence lengths

The simulated dataset’s ability to reconstruct a relationship was directly related to the length of the simulated alignment, with 500 bp performing the worst and 5000 bp performing the best (**Figure 4**). In general, the pattern showed that longer branches had greater levels of concordance. The most difficult-to-resolve clade consisted of *Conophytum uviforme, Titanopsis hugo-schlechteri, Juttadinteria attenuata, Juttadinteria kovisimontana, Dracophilus dealbatus, Psammophora modesta,* and *Ruschianthus falcatus* and was located on a branch of length 0.000696156 subs/site. The simulated genes recovered the correct relationship at 34.4% for 500 bp, 58.1% for 1000 bp, 82.2% for 2000 bp, and 98.1% for 5000 bp. The clade leading to the core Ruschioideae received 100% gCF support across all simulated gene lengths and sat on a branch of length 0.053951996.

**Figure 4.**
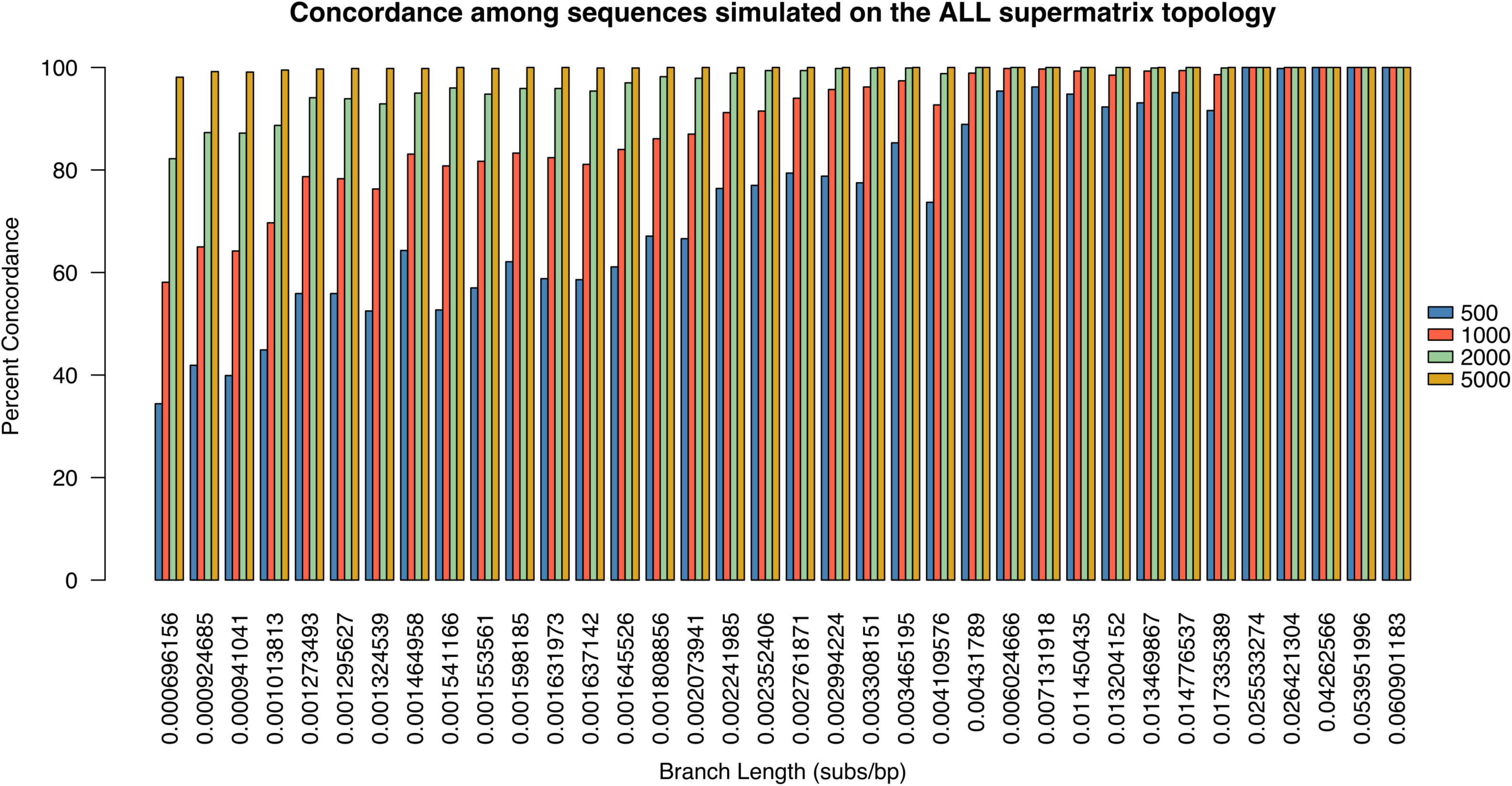
Simulated gene trees have low concordance for the short branch lengths inferred in the Ruschiodeae. The x-axis shows branch lengths of the ALL supermatrix topology and the y-axis shows the proportion of genes concordant with the species tree. Each bar color represents a different gene length corresponding to the lengths on the right.

### Gene Duplication and Functional Annotation

Gene duplication events were identified using PhyParts by examining only putative duplications at nodes with branch support ≥ 95% UFBoot among the 13,822 extracted clades. Using this cutoff, 26 nodes showed no well-supported shared gene duplications (**Supplementary Figure 7**). The node subtending the core Ruschioideae had the most gene duplications, with 52 well-supported gene duplication events across 50 homolog trees predicted at this node (**Supplementary Table 5**). Among these duplications, 10 have been reported in the literature as associated with the 3 major xeromorphic characters hypothesized to drive the rapid radiation (**Table 1**). Limitations and biases exist in this process of identifying putative duplications, and they are discussed in the Discussion.

**Table 1.** Duplicated genes at the base of Ruschioideae with potential links to key adaptations that allow for the transition to a desert environment.

| Adaptation | Panther ID | Gene Name | Putative Functional Link to Adaptation | Citation |
|---|---|---|---|---|
| Wide-Band Tracheids | PTHR11746:SF336 | O-methyltransferase family protein | Production of lignin units constituting thickened secondary cell walls in nonliving, lignified, non-perforated cells that prevent primary wall collapse under low water potential | Boerjan et al., 2003; Van Acker et al., 2013 |
| Wide-Band Tracheids | PTHR11926:SF1464 | UDP-glycosyltransferase 76B1-like / glycosyltransferase | Glycosylation of monolignol precursors into transport-competent glucoside forms for delivery to the cell wall during lignin polymerization | Morant et al., 2008; Van Acker et al., 2013 |
| Wide-Band Tracheids | PTHR19375:SF547 | 70 kDa heat shock protein (HSP70) | Chaperone activity during lignifying tissue development, linking HSP70 function to secondary cell wall biosynthesis | ul Haq et al., 2019 |
| Wide-Band Tracheids / Secondary cell wall elaboration | PTHR31444:SF8 | Protein irregular xylem 15 (IRX15) | Encodes a DUF579-domain protein essential for xylan deposition in the secondary cell wall, providing the hemicellulose scaffold into which lignin is deposited | Brown et al., 2011 |
| Flattened/cylindrical/trigonous leaf shape | PTHR47002:SF2 | Aquaporin AQPAE.A-like | Transmembrane water flux and turgor maintenance in peripheral vascular bundles surrounding water-storing tissue | Maurel et al., 2015; Afzal et al., 2016 |
| Flattened/cylindrical/trigonous leaf shape | PTHR31956:SF26 | Non-specific phospholipase C2 | Phospholipid-derived osmotic stress signaling regulating stomatal closure and cellular water status under drought | Laxalt and Munnik, 2002 |
| Flattened/cylindrical/trigonous leaf shape & Hygrochastic capsules | PTHR10791:SF22 | Bidirectional sugar transporter SWEET11 / SWEET | Photosynthate supply to achlorophyllous hydrenchyma sink tissue and Sucrose unloading from maternal vascular tissue into seed apoplast and pericarp during fruit development | Chen et al., 2012; Sosso et al., 2015; Anjali et al., 2020;<br>Pegler et al., 2023 |
| Hygrochastic capsules | PTHR45933:SF11 | Protein C2-domain ABA-related 1 | ABA-mediated regulation of seed dormancy and coat maturation governing controlled partial-release dispersal strategy | Leung & Giraudat, 1998; Finkelstein, 2013 |
| Hygrochastic capsules | PTHR45751:SF30 | E3 ubiquitin-protein ligase RGLG5 / UBC core domain-containing protein | Ubiquitin-proteasome-dependent modulation of ABA receptor and transcription factor abundance during seed maturation | Sadanandom et al., 2012; Stone, 2014 |
| Hygrochastic capsules | PTHR47979:SF56 | Ras-related protein RAB11B | Recycling endosome-based vesicle trafficking directing cell differentiation and membrane remodeling required for hygrochastic capsule opening mechanics | Rutherford & Moore, 2002; Chow et al., 2008 |

### Plastome dataset composition and phylogenetic inference

The number of reads that mapped to the plastome target file ranged from 18,108 reads for *Rivina humilis* to 34,187,292 reads for *Sesuvium portulacastrum*. The fewest plastome genes recovered were 58 for *Pleiospilos magnipunctatus,* and several samples recovered all 79 genes (**Supplementary Table 6**). The inferred relationships of the plastome phylogeny showed varying degrees of support. Those preceding the Ruschioideae were well supported (95 <= UFBOOT), while nodes within Ruschioideae, primarily along the backbone, were poorly or moderately supported (UFBOOT < 90) (**Supplementary Figure 8**). The plastome tree and the ALL supermatrix tree show a high degree of similarity, but there are several notable conflicts, primarily within the Ruschioideae clade, which were moderately supported (UFBOOT <= 95). However, higher-level relationships above the Ruschioideae clade are identical between the two trees and highly supported by the plastome data.

The most notable conflicts between the ALL supermatrix and plastome trees involve the placement of *Titanopsis, Conophytum, Ruschianthus*, and *Monilaria* (**Supplementary Figure 8**). In the ALL supermatrix tree, *Titanopsis, Conophytum*, and *Ruschianthus* are all found in a clade with [*Psammophora modesta*, *Dracophilus dealbatus* and the monophyletic *Juttadinteria*], whereas the plastome tree recovers them outside of this clade. *Monilaria* is found to be sister to the rest of Ruschioideae in the plastome tree, while the ALL supermatrix tree recovers [*Gibbaeum album, Glottiphyllum uncatum, Delosperma echinatum,* and the *Trichodiadema*] *sister to all other* Ruschieae. In addition, the plastome tree places *Gibbaeum* and *Glottiphyllum* as sister to one another, the placement of *Argyroderma* differs between trees, and the monophyly of *Stoeberia* is not supported by the plastome data (**Supplementary Figure 8**).

## Discussion

### Dense ortholog sampling was recovered despite variation in transcriptome quality

The considerable variation in BUSCO completeness scores across the 39 transcriptome assemblies most likely reflects both biological and technical factors inherent to phylotranscriptomics. Relatively low completeness in specialized succulent taxa like *Sesuvium verrucosum* (32.3%) may partly reflect divergence from the reference database rather than an ineffective assembly, a known limitation of BUSCO-based assessment in non-model organisms (**Simão et al., 2015**). Additionally, the highly succulent tissues characteristic of core Ruschioideae are known to accumulate secondary metabolites that can inhibit RNA extraction and reduce RNA yield and integrity, which could potentially contribute to lower assembly completeness in these taxa independent of assembly methodology **(Ghawana et al., 2011)**. Despite this variation, consistently high per-contig TransRate scores, particularly in nucleotide accuracy, indicate that the recovered sequences have reliable base-level accuracy for phylogeny inference. The recovery of 4,845 orthologs across at least 20 taxa shows that assembly quality was sufficient to yield a reliable, gene-rich dataset.

### Rapid radiations continue to pose challenges for systematics

To investigate relationships within the Aizoaceae, we generated nine multi-locus phylogenetic trees, eight based on filtered subsets of the nuclear data and one based on the plastome. Similar to other rapid radiations (**Cai et al., 2021**), the Aizoaceae have consistently posed a systematic challenge (**Klak et al., 2004**). Phylotranscriptomics has proven to be a powerful approach to inferring contentious relationships, and here we generated a dataset of 34 Aizoaceae to test whether this data can help resolve relationships within the family.

All nine phylogenetic hypotheses recovered identical relationships among the subfamilies, similar to other studies (**Klak et al., 2003**; **Thiede, 2004**; **Klak et al., 2007**), indicating the stability of these relationships. Within the Ruschioideae, relationships remained contentious, as in previous work (**Klak et al., 2013**; **Valente et al., 2014**). However, our supermatrix tree and coalescent tree were highly similar, except for the placement of *Titanopsis*.

The most recent study of the Ruschioideae presented a densely sampled phylogenetic tree, and despite the use of ten plastid loci, several relationships could not be reliably resolved (**Klak et al., 2013**). Notably, the study only found the *Drosanthemum* to be resolved within the Ruschioideae and most other genera formed a polytomy (**Supplementary Figure 8**). The approaches used in this paper do not directly infer polytomies, but across the Ruschioideae, short branch lengths indicative of limited information make it difficult to infer bifurcations reliably.

The percentage of genes that are concordant with species relationships is directly related to a wide range of biological factors. Decreases in gene tree concordance can indicate gene duplication and loss, hybridization, and incomplete lineage sorting (**Maddison, 1997**). In rapidly radiating groups, incomplete lineage sorting is expected to occur at higher levels because alleles have less time to fix prior to speciation. The Ruschioideae, predicted to be the fastest-radiating clade of plants (**Klak et al., 2004),** showed this exact pattern, with the lowest levels of concordance inferred among those relationships (**Figure 2**). However, the branches that led to the relationships also showed minimal information, with branch lengths as low as 0.0007 subs/site for the branch leading to *Juttadinteria attenuata, Juttadinteria kovisimontana, Dracophilus dealbatus, Psammophora modesta, Ruschianthus falcatus, Conophytum uviforme, Titanopsis hugo-schlechteri.* As such, we tested whether simulated data could infer the relationships within the Ruschioideae.

The simulations were conducted at four different lengths: 500 bp, 1000 bp, 2000 bp, and 5000 bp. Each simulation was run using the tree’s branch lengths and the general time-reversible model with gamma-rate variation. Phylogenetic trees were then inferred using the same model of evolution to create the ideal scenario for phylogenetic inference. Despite being estimated under the exact model, most 500 bp relationships were not recovered with 100% gene tree concordance with recovery rates as low as 34.4%. The use of simulated data creates an unrealistically simple scenario for tree inference, as the specified reconstruction method is identical to the simulation method. The true molecular evolutionary model under which the genes used in the study evolved is likely far more complex than the GTR model; therefore, if the relationship cannot be reconstructed in an ideal scenario, it is unlikely to be reconstructed reliably under non-ideal conditions. The 5000 bp sequences recovered relationships in 98% or more of the gene trees, indicating the importance of highly informative genes. This is likely not unexpected, as under a 500 bp simulation with a 0.0007 subs/site branch, that would be equivalent to (500*0.0007) or 0.35 subs. Under the simulation, this means roughly 35% of the genes had at least one substitution, which is roughly the gene concordance for the node. For the 5000 bp sequence, the number of substitutions would be estimated at 3.5. In a simulation experiment, length will serve as a realistic proxy for information; furthermore, the alignment length of the empirical data is not equivalent to that in simulation, as alignments will contain gaps from insertions and deletions, and therefore, an alignment of 500 bp may not have 500 characters, unlike in simulation. As real phylogenetic information is far more complex, we tested two approaches to assess the difficulty a gene tree will impose on species tree inference in the empirical data.

The first approach was to use Pythia to examine the genes for regions of likelihood space with multiple instances of the same topologies, thereby helping determine the difficulty of inferring a single tree from a sequence. In total, 2,092 genes remained after the Pythia filtration (PYTH dataset), and the inferred topology was the same as the ALL dataset. Despite recovering the same topology, the percentage of concordant genes increased at every node, and the average alignment length also increased. The majority of internal branch lengths between the two datasets decreased with filtering, indicating that a decrease in branch length did not contribute to the decrease in concordance.

The second filter was the TCA score, which was developed to infer ancient divergences (**Salichos and Rokas, 2013**). When applied to the Aizoaceae dataset, 450 genes (BIPART dataset) were recovered as being informative. This dataset again shows an increase in gene tree concordance, an increase in gene length, and a decrease in branch length compared to the ALL tree. Despite the heavy filtration and the increase in gene length compared to the PYTH dataset, the concordance was roughly equivalent between the PYTH and BIPART datasets. When the two datasets were intersected, roughly 432 genes overlapped, yielding a fourth dataset (BOTH). The fourth dataset yielded results highly similar to those of the BIPART dataset across all metrics.

Based on simulation results and filtering, it is likely that a significant portion of the difficulty in resolving these relationships stems from insufficient information. Although a lack of information can contribute to analytical errors, it also provides evidence of rapid radiation. The lack of information at the node indicates that alleles had insufficient time to accumulate substitutions sufficient to infer relationships. A biological reason for this could be reticulate evolution; however, methods that infer these processes often rely on well-resolved gene trees, and as such, this dataset will be unable to provide the necessary input for inference of reticulation.

### Gene duplication may have facilitated the radiation of Ruschioideae

Gene duplication has been shown to be a major driver of adaptations and key innovations across the tree of life (**Baker and Woollard, 2022**). Plants show an exceptional propensity for gene duplication, and adaptations such as plant carnivory (**Renner and Specht, 2012**), betalains (**Brockington et al., 2015**), and root nodulation (**Zhang et al., 2024**) have all been linked to gene duplications. Furthermore, the Asymmetric Leaves1/Rough Sheath 2/Phantastica (ARP) gene has been linked in the Aizoaceae to a duplication underlying the rapid radiation (**Illing et al., 2009**). Transcriptome datasets have proven to be a cost-effective and powerful means of inferring gene duplications and have helped increase our understanding of their prevalence and influence on shaping the tree of life. The radiation within the Aizoaceae is predicted to have been driven by a series of adaptations; as such, we examined whether gene duplications may underlie these adaptations. Of note, our analysis has two limitations that should be taken into consideration for interpretation of the results: (1) we used a strict bootstrap filter, which should help decrease false positives but may also contribute to false negatives, and (2) the tissue type was predominantly leaf tissue, which provides another potential source of missed duplications. Our goal in this exploration was to see what signal may emerge and relate to the theorized adaptations, and because we did not enter with preconceived hypotheses about which genes should show duplication, we kept this a strict, exploratory approach. As noted above, this has previously been a powerful means of identifying genes underlying adaptations and can help guide future research, but it is by no means a definitive answer or a completely inclusive list.

As with the orthologs, divergences within the Ruschioideae generally showed few well-supported gene duplications (**Supplemental Figure 5**). However, the highest duplication level occurred at the base of the Ruschioideae, which led to rapid radiation. As such, we examined the predicted functions of the 50 homologs with well-supported gene duplications at the base of the group and have further filtered the list to a series of genes that putatively underlie adaptations that have allowed the Aizoaceae to diversify in a harsh climate, and that may have been promoted by gene duplication events.

One adaptation theorized to promote diversification is wide-band tracheids, which help prevent tissue collapse during periods of low water availability. The genes identified as potentially underlying this adaptation are listed in **Table 1** and linked to the production of lignin units that form thickened secondary cell walls in non-perforated, nonliving, lignified cells found in leaves and stems, which prevent primary wall collapse during low water potential. Their functions are linked to specialized water-conducting cells in the xylem, enabling adaptations that withstand water stress. Furthermore, glycosylation of monolignol precursors into their transport-competent glucoside forms for delivery to the cell wall during lignin polymerization, and chaperone activity during lignifying tissue development, link HSP70 function to secondary cell wall biosynthesis.

Another adaptation is the flattened leaf shape with a cylindrical/trigonous shape to reduce water loss. A series of genes was identified as possibly underlying the cylindrical and trigonous succulent leaf morphology of the Ruschioideae, functioning, respectively, in transmembrane water flux and turgor maintenance, in photosynthate supply to achlorophyllous hydrenchyma sink tissue, and in phospholipid-derived osmotic stress signaling cascades that regulate stomatal closure and cellular water status under drought conditions. They have been found as strongly associated with vascular bundles that go on to become fully peripheral (PVB), forming a ring around the water-storing tissue (**Melo-de-Pinna et al., 2014**).

The third adaptation linked to the radiation of the Ruschioideae is the hygrochastic capsule, which opens in response to moisture, allowing seeds to be deposited when conditions are conducive to their success. The core Ruschioideae fruits release few seeds, ensuring seed availability during erratic rainfall, reducing secondary dispersal (limits how seeds travel), gene flow (falls close to parent plant), and promoting reproductive isolation (**Sukhorukov et al., 2023**). These genes are also reported in the literature to function, respectively, in sucrose unloading from maternal vascular tissue into the seed apoplast and the pericarp during fruit development. They are also strongly associated with ABA-mediated regulation of seed dormancy and coat maturation governing the controlled partial-release dispersal strategy, ubiquitin-proteasome-dependent modulation of ABA receptor and transcription factor abundance during seed maturation programs, and recycling endosome-based vesicle trafficking directing the specialized cell differentiation and membrane remodeling, which is required for hygrochastic capsule opening mechanics. We found that this gene encodes a DUF579-domain protein essential for xylan deposition in the secondary cell wall, which is the hemicellulose scaffold into which lignin is deposited (**Brown et al., 2011**). Since WBTs are defined by their heavily thickened, lignified secondary cell walls, the duplication of this and Protein irregular xylem 15 (IRX15) at the node subtending core Ruschioideae further supports a broader pattern of secondary cell wall elaboration coinciding with the origin of this clade. Although few genes emerge as duplicated, this adaptation is the one most likely to be influenced by the tissue type chosen.

Taken together, the dataset suggests gene duplications underlie several adaptations predicted to have facilitated movement into arid regions. Although it is difficult to provide a perfect link, this work provides a list of candidate genes that may help with future investigations into the Ruschioideae.

### Future directions for studying rapidly radiating groups

Across the tree of life, several relationships are contentious, and gene-rich datasets from transcriptomes or probes can often provide insight. These datasets, however, rely on the assumption that gene sequences have a single history, and gene sequences are rarely longer than a few thousand base pairs. One exception is the organellar genomes (**Doyle, 1992**; **Doyle, 2022**), which are typically uniparentally inherited; however, whether systematic or biological, they have shown evidence of intergenic conflict (**Goncalves et al., 2019**; **Walker et al., 2019**), as have genes themselves (**Scornavacca and Galtier, 2016**; **Mendes et al., 2019**; **Smith et al., 2020**). To address the lack of individual gene information, we examined plastome-inferred relationships, which have historically been the primary marker for Aizoaceae systematics. Given the short branch lengths, Aizoaceae and other rapidly radiating groups may yield the most reliable results from full plastome data and the use of syntenic blocks (**Rivas-González et al., 2026**), which can provide longer sequences. Although individual genes may not be able to resolve the rapid radiations, the inability to do so is, in and of itself, an important biological aspect of radiations.

Such an approach has proven pivotal in other moderately radiating groups, where transcriptome-based datasets could not fully resolve short internode branches (**Schultz et al., 2023**; **Jiang et al., 2025**). Constructing the core Ruschioideae pangenome from chromosome-level assemblies of representative taxa would enable characterization of structural genomic variation, including presence-absence and copy-number variation that transcriptome-based approaches might not capture (**Fang et al., 2024**). Additionally, comparative transcriptomic development across Ruschioideae could enable direct testing of gene duplications identified at the base of Ruschioideae and their association with tissue-specific expression divergence underlying the xeromorphic synapomorphies of this clade.

Although this study began with the aim of shedding light on contentious relationships within the Aizoaceae, it highlights the difficulties of inferring relationships in rapidly radiating groups. The short internodal branch lengths recovered across the clade suggest that additional taxon sampling will be necessary to resolve interclade relationships, and gene-dense analyses with greater sampling have shown promise for taxonomic investigations (**Kadereit et al., 2026**). As with all transcriptome-based datasets, the gene sequences recovered here reflect only the expressed portion of the genome at the time of tissue sampling, which, along with a stringent support filter, could have influenced gene duplication results. Furthermore, the association between duplicated genes in Ruschioideae and xeromorphic characters is inferred from functional annotations derived from distantly related model organisms, and direct functional validation in Aizoaceae will be required to establish causality.

## Conclusions

This work demonstrates the important role genome data will play in the future and the future directions for functional work exploring diversification in desert conditions. Although the debate over more genes versus greater taxon sampling has long been part of phylogenetics, this study shows that in datasets based on gene trees, there may be insufficient information for resolution. Transcriptomic data can help identify putative genes underlying adaptations that drive rapid radiations, but the lack of single-gene information may limit our understanding of some of the most complex regions of the tree of life.

## Acknowledgements

The authors would like to thank the Millennium Seed Bank, Cambridge University Botanic Garden and Desert Botanical Garden for providing specimens. The University of Illinois Chicago start-up funding, the University of Michigan, and the National Science Foundation (DEB 1354048) provided funding.

## Data availability

The raw-paired end sequence reads for the newly sequenced samples are available on the NCBI Sequence Read Archive (SRA) under the BioProject ID PRJNA1561618. The transcriptome assemblies and data used for the different analyses are available on Zenodo at https://doi.org/10.5281/zenodo.23772503.

## Supplementary Captions

**Supplementary Figure 1.** Results of the Basic Universal Single Copy Orthologs (BUSCO) analysis.

**Supplementary Figure 2.** TransRate2 scores divided into different error types: uniformity of the base pair calls (*sCnuc*), coverage level of the transcript (*sCov*), pairing of the mapped reads (*sCord*); and evidence of chimerism within the transcripts (*sCseg*).

**Supplementary Figure 3.** Gene trees inferred from *matK* and *rbcL* sequences extracted from the raw sequence reads with the corresponding sequences downloaded from NCBI GenBank. Sequences downloaded from GenBank have the accession ID’s next to the species names. In red are the sequences where the inferred taxa from the transcriptome are not directly sister to those downloaded from NCBI.

**Supplementary Figure 4.** Number of sequences in each inferred homolog cluster by round. We removed any cluster that did not change in size during the cleaning procedure (Blue Squares), and we continued cleaning any that did (Red Circles).

**Supplementary Figure 5.** The species relationships inferred by each dataset and for each method. Support values on the Supermatrix trees are in Ultrafast Bootstrap values, and those on the coalescent-based (Astral) trees are in Local Posterior Probabilities.

**Supplementary Figure 6.** The Supermatrix tree inferred from the ALL dataset with the 2,857 gene trees that could be rooted on the outgroups mapped to it. Pie charts depict concordance, conflict, and lack of information at each node.

**Supplementary Figure 7.** The Supermatrix tree inferred from the ALL dataset, with the number of inferred duplications at each node. A) The total number of duplications inferred to have occurred at the node. B) The total number of homolog trees with an inferred duplication at that node.

**Supplementary Figure 8.** The relationships inferred using the complete plastome supermatrix and the hypothesized relationships from Klak et al., 2013.

**Supplementary Table 1.** The samples newly obtained for this study, with voucher information, location of the sample in living collections, and previous groupings.

**Supplementary Table 2.** The total number of paired-end reads for each sample and the SRA Accession for samples.

**Supplementary Table 3.** The composition of the bait file used to extract plastome reads and the accession number for each of the plastomes downloaded for the bait file.

**Supplementary Table 4.** The number of contigs and length of contigs used to test whether multiple copies exist within a single set of reads.

**Supplementary Table 5.** Gene duplications and predicted functions for homolog clusters inferred to have duplicated at the base of the Ruschioideae.

**Supplementary Table 6.** The number of reads and number of predicted genes for the plastomes of the species in the study.

