## Supplementary Figure 1 for "Limitations of gene-dense datasets for inferring relationships in rapid radiations: An example from the Ice Plants"

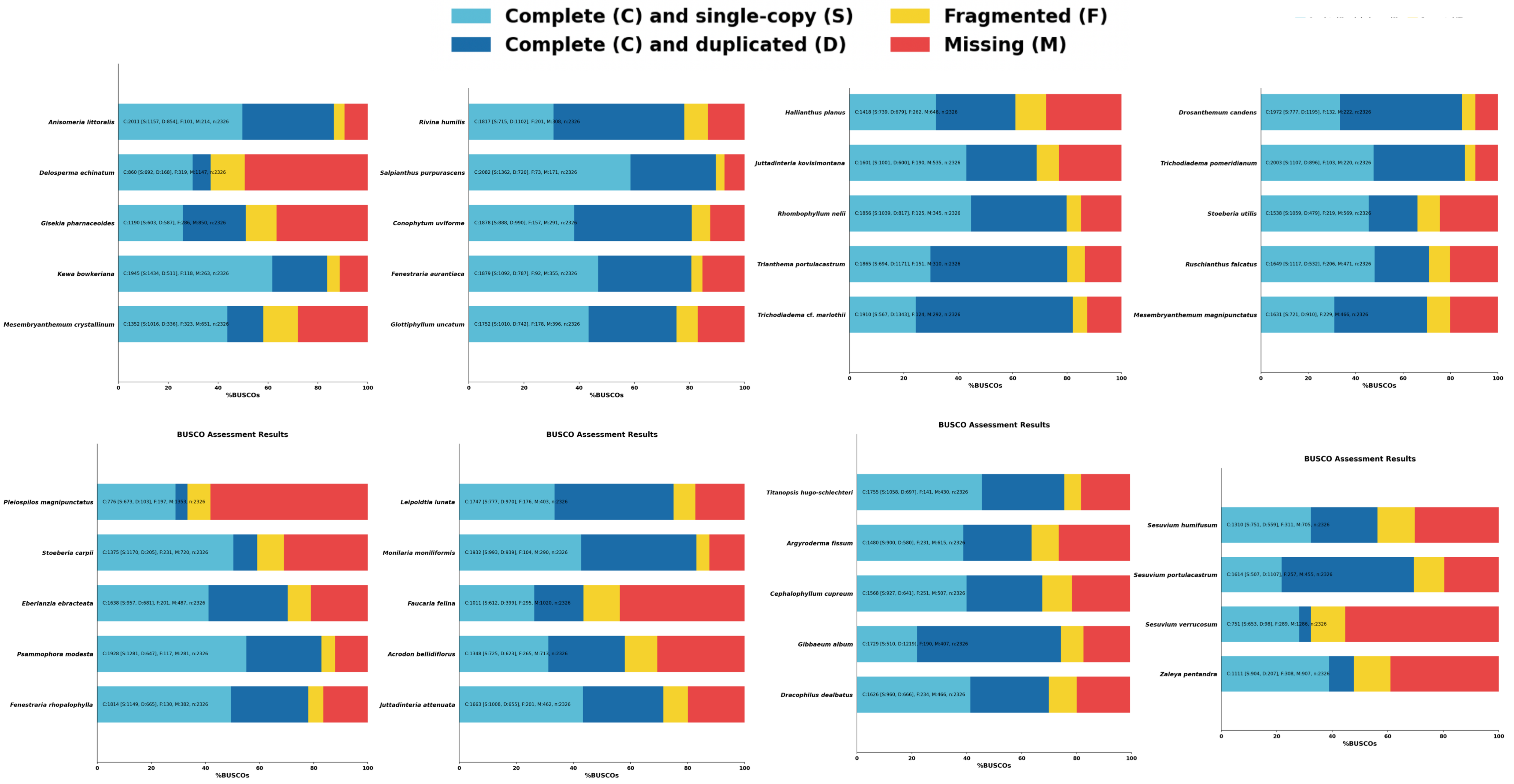

Complete (C) and single-copy (S)
Complete (C) and duplicated (D)
Fragmented (F)
Missing (M)
Anisomeria littoralis
C:2011 [S:1157, D:854], F:101, M:214, n:2326
Delosperma echinatum
C:860 [S:692, D:168], F:319, M:1147, n:2326
Gisekia pharnaceoides
C:1190 [S:603, D:587], F:286, M:850, n:2326
Kewa bowkeriana
C:1945 [S:1434, D:511], F:118, M:263, n:2326
Mesembryanthemum crystallinum
C:1352 [S:1016, D:336], F:323, M:651, n:2326
%BUSCOs
Rivina humilis
C:1817 [S:715, D:1102], F:201, M:308, n:2326
Salpianthus purpurascens
C:2082 [S:1362, D:720], F:73, M:171, n:2326
Conophytum uviforme
C:1878 [S:888, D:990], F:157, M:291, n:2326
Fenestraria aurantiaca
C:1879 [S:1092, D:787], F:92, M:355, n:2326
Glottiphyllum uncatum
C:1752 [S:1010, D:742], F:178, M:396, n:2326
%BUSCOs
Hallianthus planus
C:1418 [S:739, D:679], F:262, M:646, n:2326
Juttadinteria kovisimontana
C:1601 [S:1001, D:600], F:190, M:535, n:2326
Rhombophyllum nelii
C:1856 [S:1039, D:817], F:125, M:345, n:2326
Trianthema portulacastrum
C:1865 [S:694, D:1171], F:151, M:310, n:2326
Trichodiadema cf. marlothii
C:1910 [S:567, D:1343], F:124, M:292, n:2326
%BUSCOs
Drosanthemum candens
C:1972 [S:777, D:1195], F:132, M:222, n:2326
Trichodiadema pomeridianum
C:2003 [S:1107, D:896], F:103, M:220, n:2326
Stoeberia utilis
C:1538 [S:1059, D:479], F:219, M:569, n:2326
Ruschianthus falcatus
C:1649 [S:1117, D:532], F:206, M:471, n:2326
Mesembryanthemum magnipunctatus
C:1631 [S:721, D:910], F:229, M:466, n:2326
%BUSCOs
BUSCO Assessment Results
Pleiospilos magnipunctatus
C:776 [S:673, D:103], F:197, M:1353, n:2326
Stoeberia carpii
C:1375 [S:1170, D:205], F:231, M:720, n:2326
Eberlanzia ebracteata
C:1638 [S:957, D:681], F:201, M:487, n:2326
Psammophora modesta
C:1928 [S:1281, D:647], F:117, M:281, n:2326
Fenestraria rhopalophylla
C:1814 [S:1149, D:665], F:130, M:382, n:2326
%BUSCOs
BUSCO Assessment Results
Leipoldtia lunata
C:1747 [S:777, D:970], F:176, M:403, n:2326
Monilaria moniliformis
C:1932 [S:993, D:939], F:104, M:290, n:2326
Faucaria felina
C:1011 [S:612, D:399], F:295, M:1020, n:2326
Acrodon bellidiflorus
C:1348 [S:725, D:623], F:265, M:713, n:2326
Juttadinteria attenuata
C:1663 [S:1008, D:655], F:201, M:462, n:2326
%BUSCOs
BUSCO Assessment Results
Titanopsis hugo-schlechteri
C:1755 [S:1058, D:697], F:141, M:430, n:2326
Argyroderma fissum
C:1480 [S:900, D:580], F:231, M:615, n:2326
Cephalophyllum cupreum
C:1568 [S:927, D:641], F:251, M:507, n:2326
Gibbaeum album
C:1729 [S:510, D:1219], F:190, M:407, n:2326
Dracophilus dealbatus
C:1626 [S:960, D:666], F:234, M:466, n:2326
%BUSCOs
BUSCO Assessment Results
Sesuvium humifusum
C:1310 [S:751, D:559], F:311, M:705, n:2326
Sesuvium portulacastrum
C:1614 [S:507, D:1107], F:257, M:455, n:2326
Sesuvium verrucosum
C:751 [S:653, D:98], F:289, M:1286, n:2326
Zaleya pentandra
C:1111 [S:904, D:207], F:308, M:907, n:2326
%BUSCOs
