## Supplementary figures and images for "Limitations of gene-dense datasets for inferring relationships in rapid radiations: An example from the Ice Plants"

### Supplementary Figure 2

#### TransRate Assembly Scores

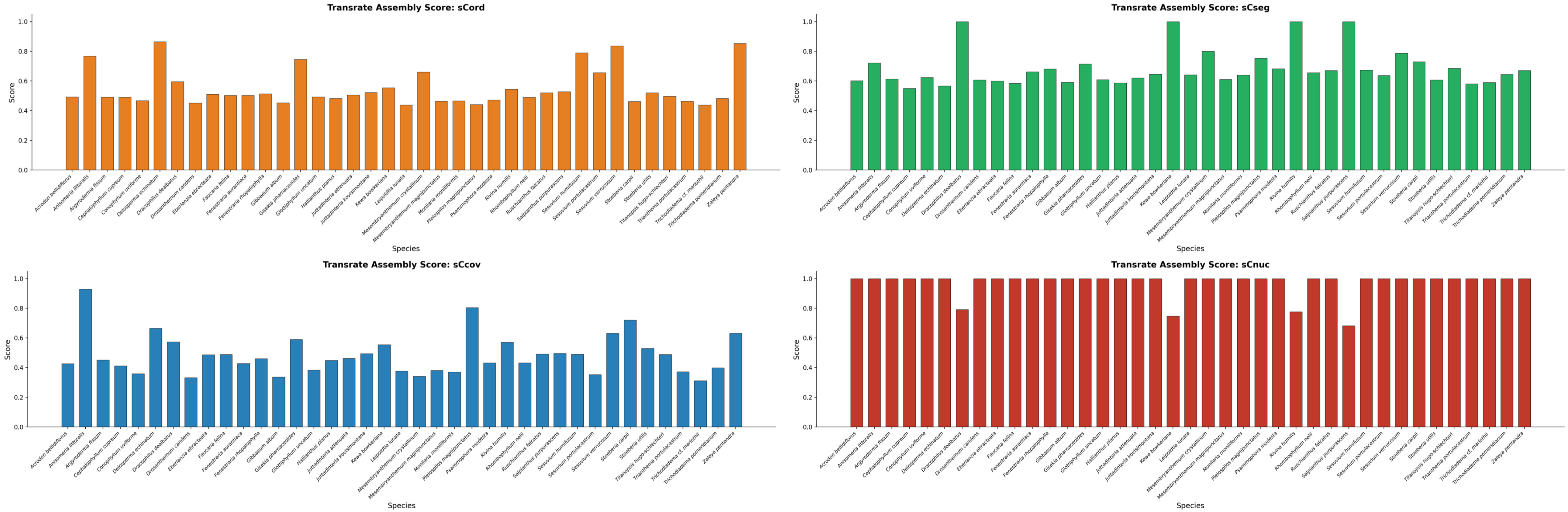

### Supplementary Figure 5

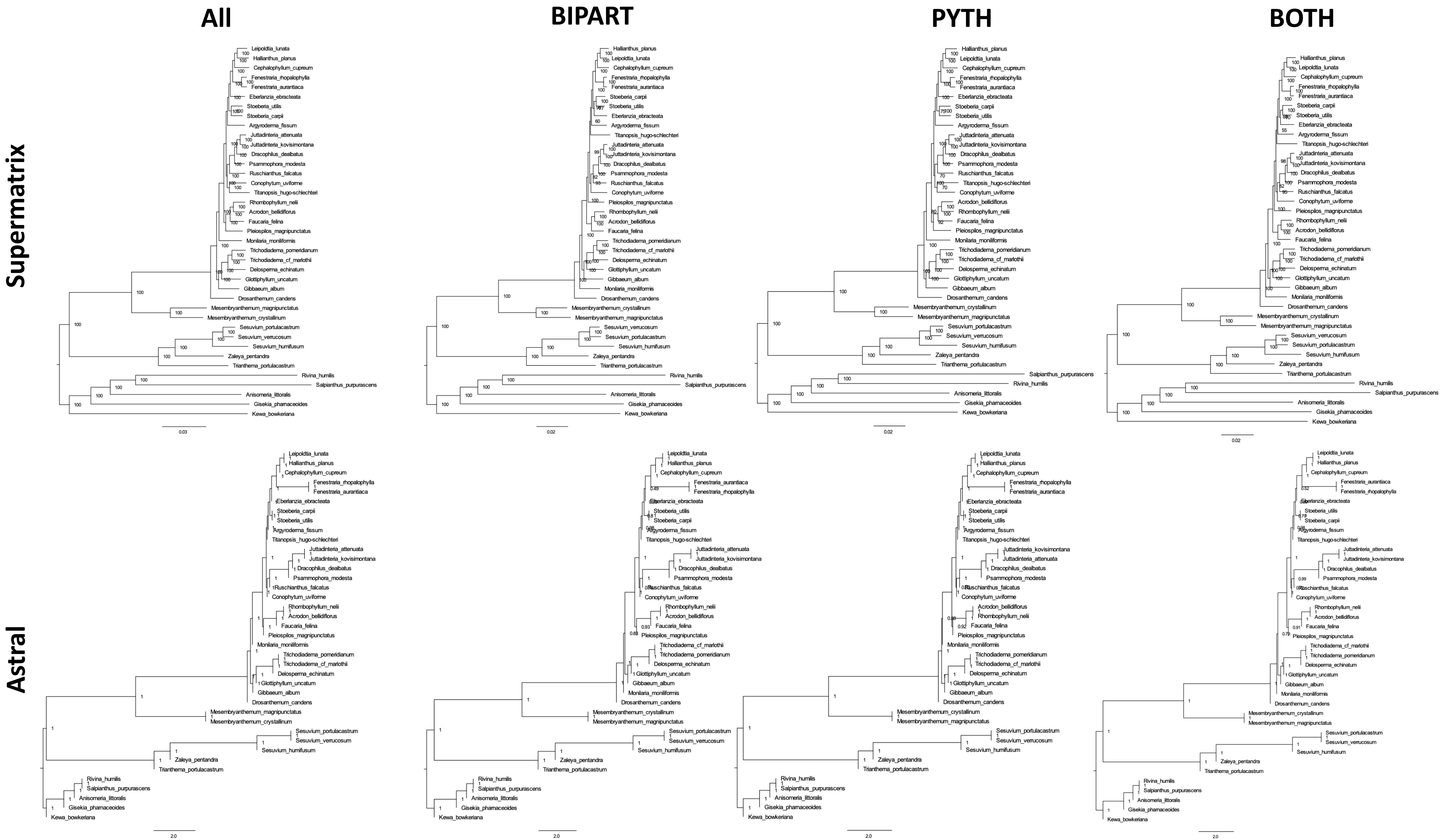
All
BIPART
PYTH
BOTH
Supermatrix
Astral
0.03
0.02
0.02
0.02
2.0
2.0
2.0
2.0

### Supplementary Figure 6

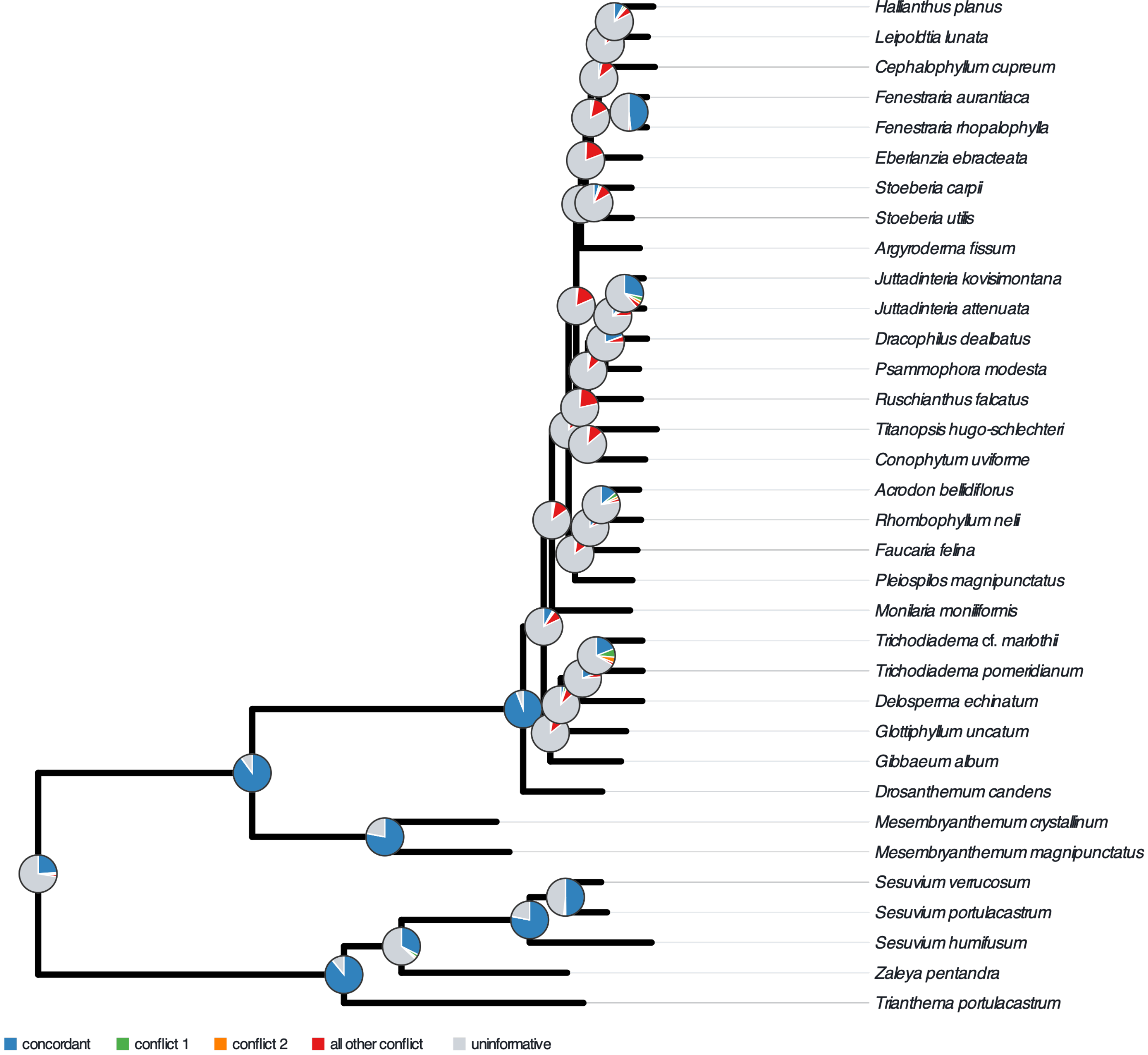


2857 gene trees · 39 taxa · gene tree branches with support < 95 collapsed

### Supplementary Figure 8

##### Plastome Tree

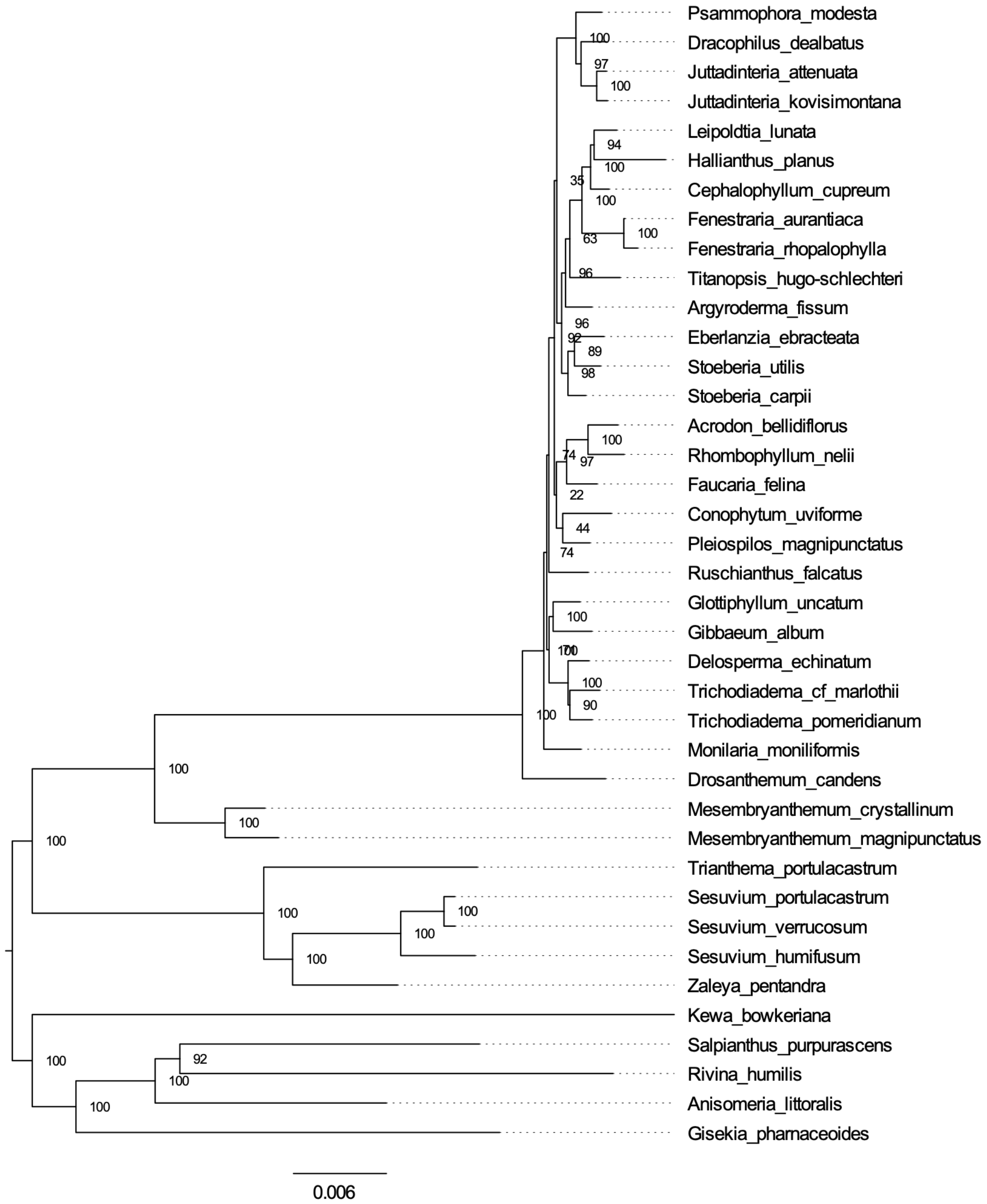


##### Klak et al., 2013

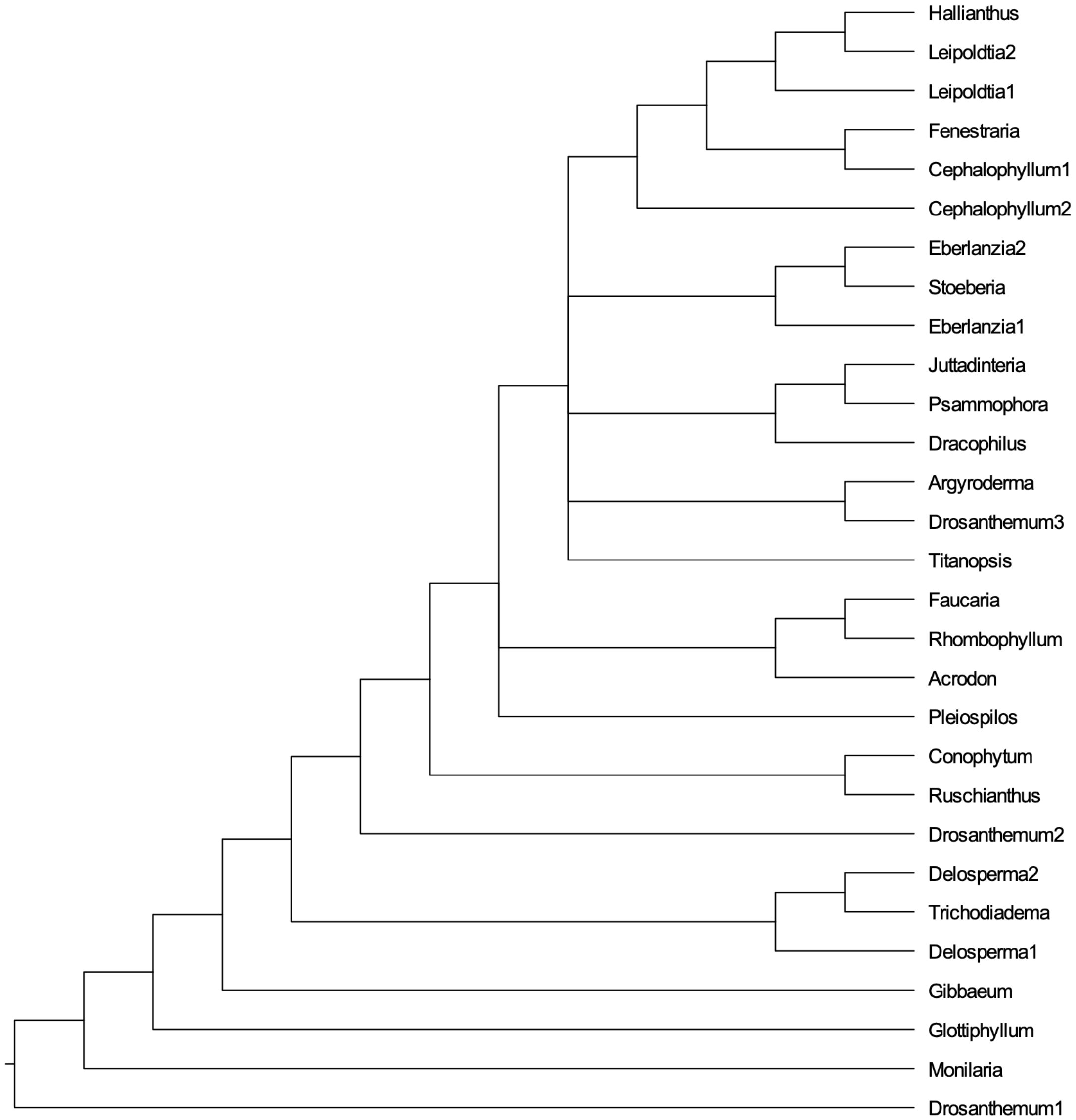
