## Supplementary Figure 3 for "Limitations of gene-dense datasets for inferring relationships in rapid radiations: An example from the Ice Plants"

*matK*

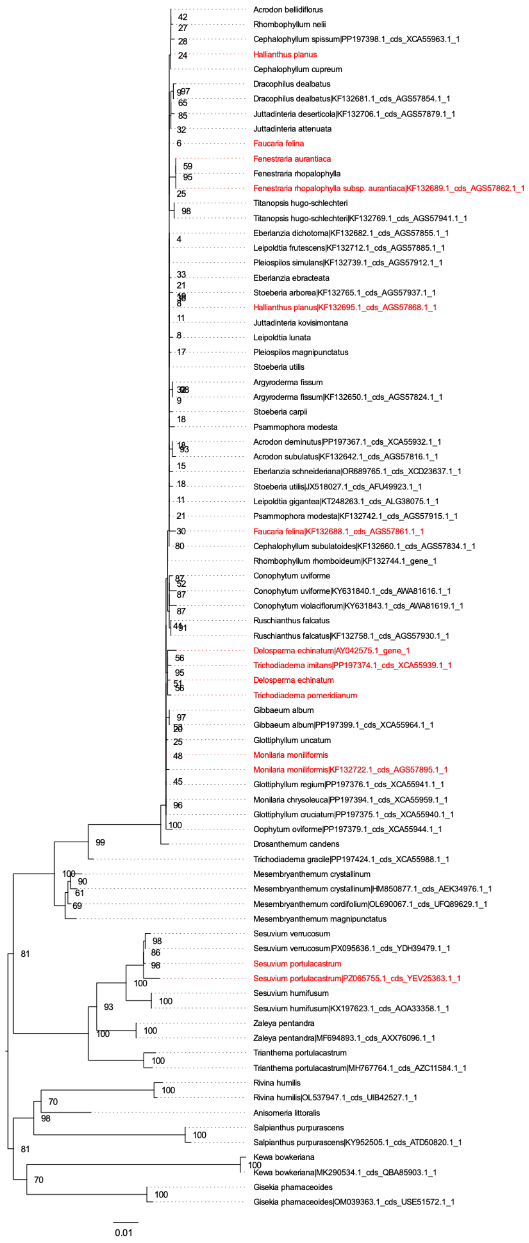
Acrodon bellidiflorus
Rhombophyllum nelii
Cephalophyllum spissum|PP197398.1_cds_XCA55963.1_1
Hallianthus planus
Cephalophyllum cupreum
Dracophilus dealbatus
Dracophilus dealbatus|KF132681.1_cds_AGS57854.1_1
Juttadinteria deserticola|KF132706.1_cds_AGS57879.1_1
Juttadinteria attenuata
Faucaria felina
Fenestraria aurantiaca
Fenestraria rhopalophylla
Fenestraria rhopalophylla subsp. aurantiaca|KF132689.1_cds_AGS57862.1_1
Titanopsis hugo-schlechteri
Titanopsis hugo-schlechteri|KF132769.1_cds_AGS57941.1_1
Eberlanzia dichotoma|KF132682.1_cds_AGS57855.1_1
Leipoldtia frutescens|KF132712.1_cds_AGS57885.1_1
Pleiospilos simulans|KF132739.1_cds_AGS57912.1_1
Eberlanzia ebracteata
Stoeberia arborea|KF132765.1_cds_AGS57937.1_1
Hallianthus planus|KF132695.1_cds_AGS57868.1_1
Juttadinteria kovisimontana
Leipoldtia lunata
Pleiospilos magnipunctatus
Stoeberia utilis
Argyroderma fissum
Argyroderma fissum|KF132650.1_cds_AGS57824.1_1
Stoeberia carpii
Psammophora modesta
Acrodon deminutus|PP197367.1_cds_XCA55932.1_1
Acrodon subulatus|KF132642.1_cds_AGS57816.1_1
Eberlanzia schneideriana|OR689765.1_cds_XCD23637.1_1
Stoeberia utilis|JX518027.1_cds_AFU49923.1_1
Leipoldtia gigantea|KT248263.1_cds_ALG38075.1_1
Psammophora modesta|KF132742.1_cds_AGS57915.1_1
Faucaria felina|KF132688.1_cds_AGS57861.1_1
Cephalophyllum subulatoides|KF132660.1_cds_AGS57834.1_1
Rhombophyllum rhomboideum|KF132744.1_gene_1
Conophytum uviforme
Conophytum uviforme|KY631840.1_cds_AWA81616.1_1
Conophytum violaciflorum|KY631843.1_cds_AWA81619.1_1
Ruschianthus falcatus
Ruschianthus falcatus|KF132758.1_cds_AGS57930.1_1
Delosperma echinatum|AY042575.1_gene_1
Trichodiadema imitans|PP197374.1_cds_XCA55939.1_1
Delosperma echinatum
Trichodiadema pomeridianum
Gibbaeum album
Gibbaeum album|PP197399.1_cds_XCA55964.1_1
Glottiphyllum uncatum
Monilaria moniliformis
Monilaria moniliformis|KF132722.1_cds_AGS57895.1_1
Glottiphyllum regium|PP197376.1_cds_XCA55941.1_1
Monilaria chrysoleuca|PP197394.1_cds_XCA55959.1_1
Glottiphyllum cruciatum|PP197375.1_cds_XCA55940.1_1
Oophytum oviforme|PP197379.1_cds_XCA55944.1_1
Drosanthemum candens
Trichodiadema gracile|PP197424.1_cds_XCA55988.1_1
Mesembryanthemum crystallinum
Mesembryanthemum crystallinum|HM850877.1_cds_AEK34976.1_1
Mesembryanthemum cordifolium|OL690067.1_cds_UFQ89629.1_1
Mesembryanthemum magnipunctatus
Sesuvium verrucosum
Sesuvium verrucosum|PX095636.1_cds_YDH39479.1_1
Sesuvium portulacastrum
Sesuvium portulacastrum|PZ065755.1_cds_YEV25363.1_1
Sesuvium humifusum
Sesuvium humifusum|KX197623.1_cds_AOA33358.1_1
Zaleya pentandra
Zaleya pentandra|MF694893.1_cds_AXX76096.1_1
Trianthema portulacastrum
Trianthema portulacastrum|MH767764.1_cds_AZC11584.1_1
Rivina humilis
Rivina humilis|OL537947.1_cds_UIB42527.1_1
Anisomeria littoralis
Salpianthus purpurascens
Salpianthus purpurascens|KY952505.1_cds_ATD50820.1_1
Kewa bowkeriana
Kewa bowkeriana|MK290534.1_cds_QBA85903.1_1
Gisekia pharnaceoides
Gisekia pharnaceoides|OM039363.1_cds_USE51572.1_1
0.01

*rbcL*

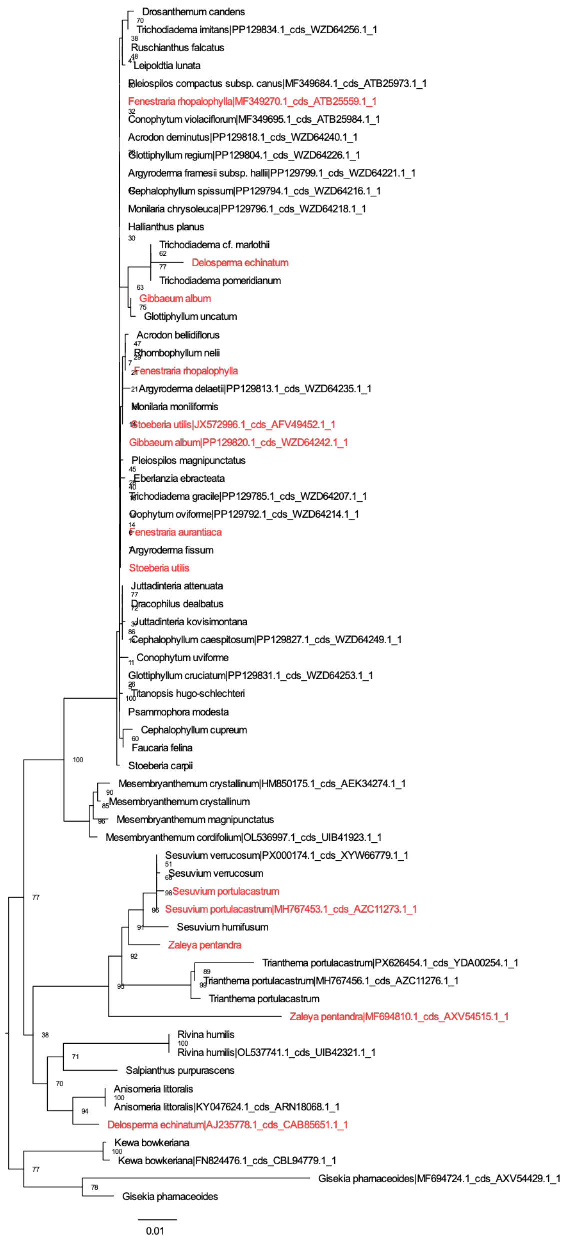
Drosanthemum candens
Trichodiadema imitans|PP129834.1_cds_WZD64256.1_1
Ruschianthus falcatus
Leipoldtia lunata
Pleiospilos compactus subsp. canus|MF349684.1_cds_ATB25973.1_1
Fenestraria rhopalophylla|MF349270.1_cds_ATB25559.1_1
Conophytum violaciflorum|MF349695.1_cds_ATB25984.1_1
Acrodon deminutus|PP129818.1_cds_WZD64240.1_1
Glottiphyllum regium|PP129804.1_cds_WZD64226.1_1
Argyroderma framesii subsp. hallii|PP129799.1_cds_WZD64221.1_1
Cephalophyllum spissum|PP129794.1_cds_WZD64216.1_1
Monilaria chrysoleuca|PP129796.1_cds_WZD64218.1_1
Hallianthus planus
Trichodiadema cf. marlothii
Delosperma echinatum
Trichodiadema pomeridianum
Gibbaeum album
Glottiphyllum uncatum
Acrodon bellidiflorus
Rhombophyllum nelii
Fenestraria rhopalophylla
Argyroderma delaetii|PP129813.1_cds_WZD64235.1_1
Monilaria moniliformis
Stoeberia utilis|JX572996.1_cds_AFV49452.1_1
Gibbaeum album|PP129820.1_cds_WZD64242.1_1
Pleiospilos magnipunctatus
Eberlanzia ebracteata
Trichodiadema gracile|PP129785.1_cds_WZD64207.1_1
Oophytum oviforme|PP129792.1_cds_WZD64214.1_1
Fenestraria aurantiaca
Argyroderma fissum
Stoeberia utilis
Juttadinteria attenuata
Dracophilus dealbatus
Juttadinteria kovisimontana
Cephalophyllum caespitosum|PP129827.1_cds_WZD64249.1_1
Conophytum uviforme
Glottiphyllum cruciatum|PP129831.1_cds_WZD64253.1_1
Titanopsis hugo-schlechteri
Psammophora modesta
Cephalophyllum cupreum
Faucaria felina
Stoeberia carpii
Mesembryanthemum crystallinum|HM850175.1_cds_AEK34274.1_1
Mesembryanthemum crystallinum
Mesembryanthemum magnipunctatus
Mesembryanthemum cordifolium|OL536997.1_cds_UIB41923.1_1
Sesuvium verrucosum|PX000174.1_cds_XYW66779.1_1
Sesuvium verrucosum
Sesuvium portulacastrum
Sesuvium portulacastrum|MH767453.1_cds_AZC11273.1_1
Sesuvium humifusum
Zaleya pentandra
Trianthema portulacastrum|PX626454.1_cds_YDA00254.1_1
Trianthema portulacastrum|MH767456.1_cds_AZC11276.1_1
Trianthema portulacastrum
Zaleya pentandra|MF694810.1_cds_AXV54515.1_1
Rivina humilis
Rivina humilis|OL537741.1_cds_UIB42321.1_1
Salpianthus purpurascens
Anisomeria littoralis
Anisomeria littoralis|KY047624.1_cds_ARN18068.1_1
Delosperma echinatum|AJ235778.1_cds_CAB85651.1_1
Kewa bowkeriana
Kewa bowkeriana|FN824476.1_cds_CBL94779.1_1
Gisekia pharnaceoides|MF694724.1_cds_AXV54429.1_1
Gisekia pharnaceoides
0.01
