## Supplementary Figure 4 for "Limitations of gene-dense datasets for inferring relationships in rapid radiations: An example from the Ice Plants"

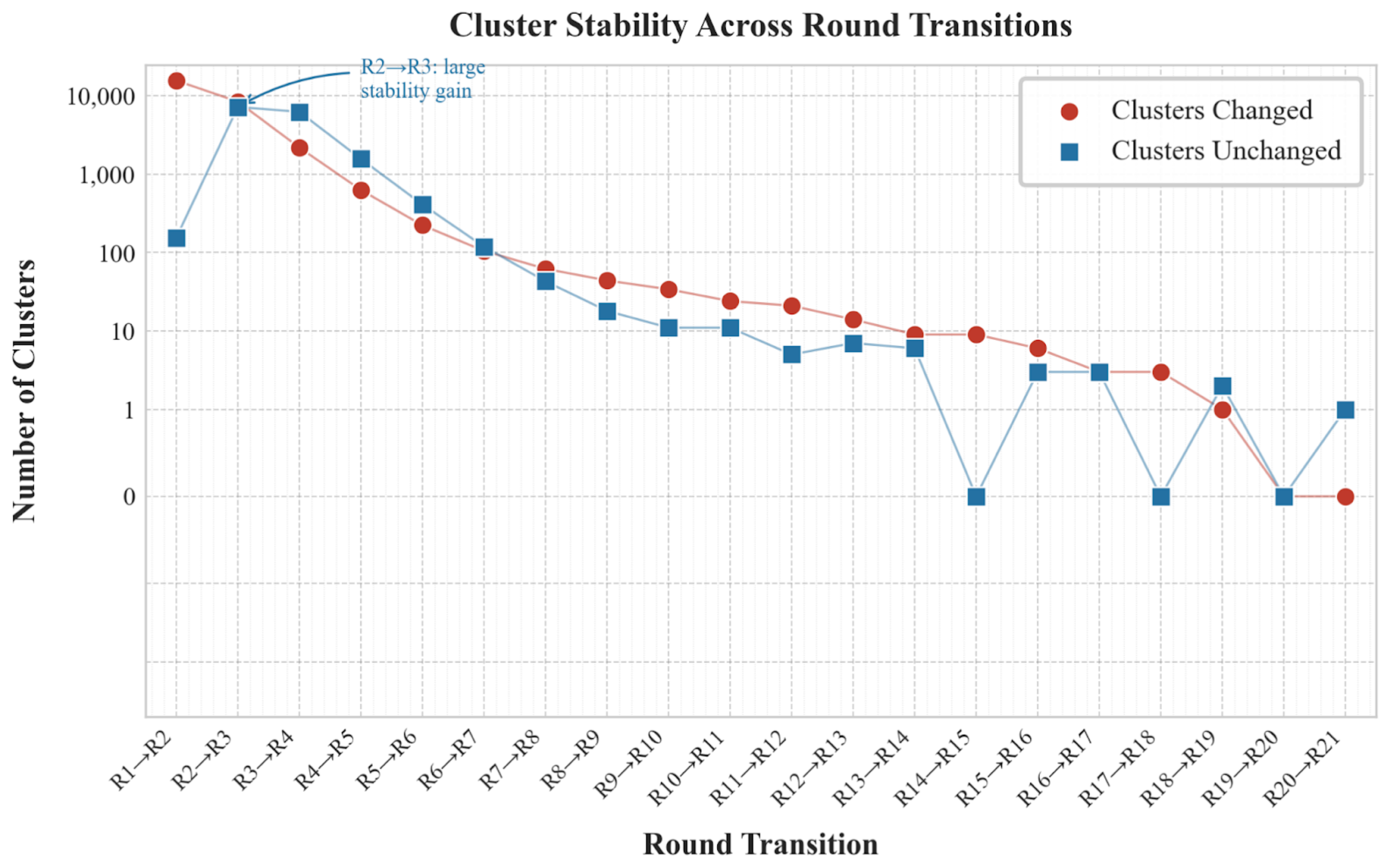

Cluster Stability Across Round Transitions
R2→R3: large stability gain
Clusters Changed
Clusters Unchanged
Number of Clusters
10,000
1,000
100
10
1
0
R1→R2
R2→R3
R3→R4
R4→R5
R5→R6
R6→R7
R7→R8
R8→R9
R9→R10
R10→R11
R11→R12
R12→R13
R13→R14
R14→R15
R15→R16
R16→R17
R17→R18
R18→R19
R19→R20
R20→R21
Round Transition
