## Supplementary Figure 7 for "Limitations of gene-dense datasets for inferring relationships in rapid radiations: An example from the Ice Plants"

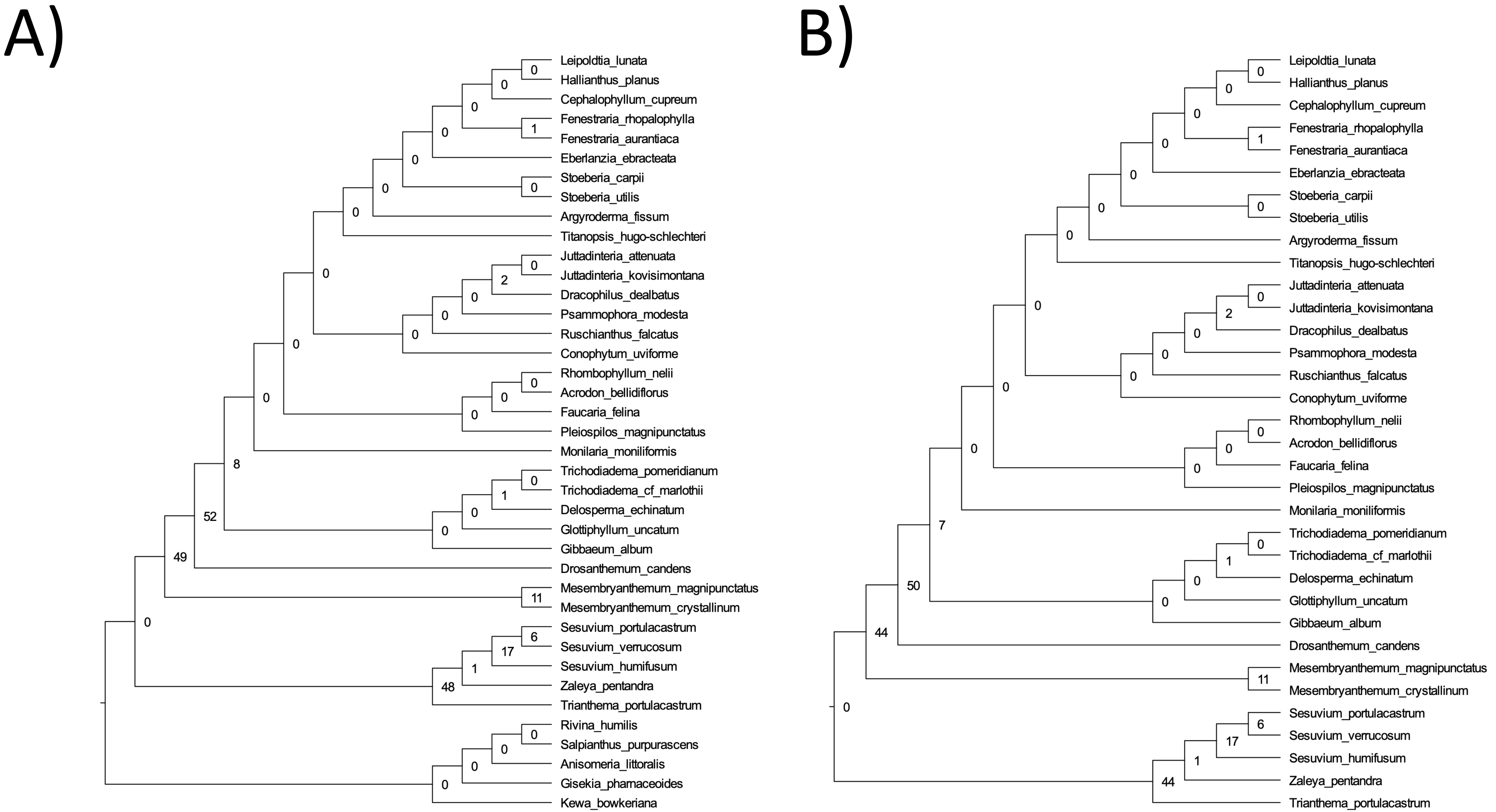
A)
B)
Leipoldtia_lunata
Hallianthus_planus
Cephalophyllum_cupreum
Fenestraria_rhopalophylla
Fenestraria_aurantiaca
Eberlanzia_ebracteata
Stoeberia_carpii
Stoeberia_utilis
Argyroderma_fissum
Titanopsis_hugo-schlechteri
Juttadinteria_attenuata
Juttadinteria_kovisimontana
Dracophilus_dealbatus
Psammophora_modesta
Ruschianthus_falcatus
Conophytum_uviforme
Rhombophyllum_nelii
Acrodon_bellidiflorus
Faucaria_felina
Pleiospilos_magnipunctatus
Monilaria_moniliformis
Trichodiadema_pomeridianum
Trichodiadema_cf_marlothii
Delosperma_echinatum
Glottiphyllum_uncatum
Gibbaeum_album
Drosanthemum_candens
Mesembryanthemum_magnipunctatus
Mesembryanthemum_crystallinum
Sesuvium_portulacastrum
Sesuvium_verrucosum
Sesuvium_humifusum
Zaleya_pentandra
Trianthema_portulacastrum
Rivina_humilis
Salpianthus_purpurascens
Anisomeria_littoralis
Gisekia_pharnaceoides
Kewa_bowkeriana
